# ANS: Adjusted Neighborhood Scoring to improve gene signature-based cell annotation in single-cell RNA-seq data

**DOI:** 10.1101/2023.09.20.558114

**Authors:** Laure Ciernik, Agnieszka Kraft, Florian Barkmann, Josephine Yates, Valentina Boeva

**Affiliations:** ETH Zurich, Department of Computer Science, Institute for Machine Learning, 8092 Zurich, Switzerland; Machine Learning Group, Technische Universität Berlin, 10587 Berlin, Germany; Berlin Institute for the Foundations of Learning and Data (BIFOLD), 10587 Berlin, Germany; Hector Fellow Academy, 76131 Karlsruhe, Germany; ETH AI Center, ETH Zürich, Zurich Switzerland; INSERM, U1016, Cochin Institute, CNRS UMR8104, Paris Descartes University, Paris, France; Swiss Institute of Bioinformatics (SIB), Zurich, Switzerland; University Hospital Zurich, Department of Thoracic Surgery, 8092 Zurich

**Keywords:** scRNA-seq, gene signatures, scoring methods, cell type annotation, cell state annotation

## Abstract

In the field of single-cell RNA sequencing (scRNA-seq), gene signature scoring is integral for pinpointing and characterizing distinct cell populations. However, challenges arise in ensuring the robustness and comparability of scores across various gene signatures and across different batches and conditions. Here, we evaluated the stability of established methods such as Scanpy, UCell, and JASMINE in the context of scoring cells of different types and states. On eight cancer and healthy scRNA-seq datasets, we reported that none of the existing methods provide fair gene signature scores that can be used in unsupervised cell state annotation based on the highest signature values. Addressing this challenge, we introduced a new scoring method, the Adjusted Neighbourhood Scoring (ANS), that builds on the traditional Scanpy method and improves the handling of the control gene sets. We further exemplified the usability of ANS scoring in differentiating between cancer-associated fibroblasts and malignant cells undergoing epithelial-mesenchymal transition (EMT) in four cancer types, and evidenced excellent classification performance (AUCPR train: 0.95-0.99, AUCPR test: 0.91-0.99). In summary, our research introduces ANS as a robust and deterministic scoring approach that enables the comparison of diverse gene signatures and score-based annotation of cell types and states. The results of our study contribute to the development of more accurate and reliable methods for analyzing scRNA-seq data.

## Introduction

High-throughput single-cell RNA sequencing (scRNA-seq) is a powerful technology to profile the transcriptome at the cellular level^1^. Gene expression from single cells allows for quantifying cell types and states, analyzing inter- and intra-sample heterogeneity, discovering cell differentiation trajectories, and constructing gene regulatory networks^2^. The interpretation of scRNA-seq data can be challenging due to high dimensionality, batch effects, dropout, and transcriptional noise^3^. To work reliably on scRNA-seq measurements, the methods must address the inherent variability and noise in these data. This is especially important when evaluating cell states and programs through gene signature scoring. Thus, robust scoring tools are essential for the reliable characterization of cell states and programs.

Gene signature scoring on scRNA-seq data measures the activity of biological processes and transcriptional cell states for each cell profiled. Gene signatures are sets of genes associated with a specific biological signal, transcriptional state, cell type, or cell state and are used as a surrogate representation for a biological phenotype^4^. During gene signature scoring, the information in the high-dimensional gene expression data is condensed into a per-cell score, which can then be used for downstream analysis. One key application is cell annotation as it offers a highly efficient and reliable method for classifying cells into types and states^5^. By ranking the scores for potential cell types and states, a higher score indicates a stronger match to a specific cell type’s phenotype. Cells can then be labelled using various strategies, such as setting a score threshold, selecting the highest score, or comparing the relative difference between scores. Notably, the quality of gene signatures plays a critical role in this process, as the accuracy of the score depends on how well the selected genes reflect the underlying biological process, regardless of the performance of the scoring method itself.

Numerous techniques for gene signature scoring in bulk RNA-seq and scRNA-seq have been developed lately. Recent studies, however, showed that signature scoring methods created for bulk RNA-seq, such as ssGSEA^6^ and GSVA^7^, were not fit for scRNA-seq data since they were more susceptible to dropouts, and suffered from imbalanced expressions of genes in cancer cells versus non-malignant cells in tumor samples^8^. Therefore, researchers were advised to use single cell-specific methods, such as signature scoring methods of Scanpy^9,10^, Seurat^11,12^, UCell^13^, and JASMINE (Jointly Assessing Signature Mean and Inferring Enrichment)^8^. Gibbs et al. propose an alternative approach to address the challenges of applying scoring methods designed for bulk data to single-cell datasets. They introduce GSSNNG, a method which overcomes sparsity and technical noise in single-cell data by using a nearest neighbor graph of cells for gene expression matrix smoothing before gene signature scoring^14^. While smoothing reduces noise, it is important to note the trade-off between removing variance and introducing bias in single-cell studies^15,16^. Additionally, this method requires the user to define cell groupings, which can be challenging and may significantly impact the resulting neighbor graphs.

The scoring methods implemented in the scRNA-seq analysis packages Scanpy^9,10^ and Seurat^11,12^, are based on the procedure described by Tirosh *et al.*^17^, which computes scores by averaging the difference in expression of the signature and control genes. Due to slight implementation differences, we distinguish between the Scanpy and the Seurat methods in this study (Methods). Although widely used, these scoring approaches may show some drawbacks. As in Scanpy and Seurat the control genes are selected by binning genes based on comparable mean expression, the scores may be biased by the number of bins and the behavior of the mean expression curve.

Two other popular scoring approaches, UCell^13^, which is an extension of AUCell^18^, and JASMINE^8^, use rank statistics to increase the stability of results in case of strong technical variation and batch effects. UCell is solely based on Mann-Whitney U statistics^13^. It is expected to be robust against technical variation as it only uses per-cell gene rank information. JASMINE computes scores by averaging the mean of signature gene ranks and an enrichment value^8^. The enrichment values can correspond to odds ratios or the likelihood, referred to as OR and LH in our benchmarks.

Although Noureen et al.^8^ investigated the robustness of various signature scoring methods for bulk and single-cell RNA-seq, focusing on sensitivity and specificity across *in silico* datasets, the study did not address score range comparability. In contrast, Wang and Thakar^19^ provided a broad comparison of several methods, including UCell, AUCell, JASMINE, and Seurat scoring, analyzing their performance under different factors, such as cell count, gene set size, noise, condition-specific genes, and zero imputation. However, their study did not include cancer datasets or evaluate these methods for score-based cell labeling. Therefore, a novel benchmark is needed to interrogate the robustness of gene signature scoring methods under the variation in sample cell composition and batch effects, ensuring their reliability and utility for score-based annotation of cell types and states in downstream analysis.

In this work, we benchmark and analyze the stability of gene signature scores provided by the most popular cell scoring methods, Scanpy, Seurat, UCell, and JASMINE. Using eight human healthy and cancer scRNA-seq datasets, we find that accurate unsupervised cell state annotation using gene signature scores is not possible based on scores calculated by the benchmarked methods, due to biased score ranges returned by all methods tested. We present an improvement of Tirosh’s scoring method, Adjusted Neighbourhood Scoring (ANS), which is more resistant to single-cell technology biases than the original approach. We show that ANS is robust with regard to most influencing factors, and returns comparable scores for multiple signatures that can be used for accurate cell type and cell state annotation. To showcase the usability of ANS to differentiate cells with related gene signatures, we use our method to devise a signature for mesenchymal-like cancer cells that discriminates mesenchymal-like malignant cells from tumor-infiltrating fibroblasts across different cancer types. Overall, our improvement to signature scoring methods, generating accurate and comparable gene signature scores, enables more reliable cell type and cell state annotation in single-cell data across diverse cell types and conditions.

## Results

### Adjusted Neighbourhood Scoring (ANS) method

We proposed to improve the widely used gene signature scoring method suggested by Tirosh *et al.*^17^, which had been implemented in Seurat^11^, and with minor modifications in Scanpy^9^. In brief, the Tirosh method computes the gene signature scores for each cell by subtracting a control expression from the signature genes’ expressions. The major drawback of the Tirosh signature scoring method comes from the procedure of choosing control genes using fixed binning of genes according to their average gene expression (default: 25 bins, Suppl. Figure S1). As verified by our benchmarking, this procedure may create artifacts for the estimates of expression gain, specifically for highly expressed genes frequently used in gene signatures. Our proposed method, called Adjusted Neighborhood scoring (ANS), solved this issue by selecting control gene sets separately for each signature gene so that the averaged expression of the control gene set and the signature gene matches (Methods). The ANS method was implemented in Python and R (package https://github.com/lciernik/ANS_signature_scoring).

### Design of the benchmark

We benchmarked ANS and other gene signature scoring methods, including the original Tirosh method, assessed the stability of the signature scores, and highlighted the importance of the choice of control genes. We compared eight gene signature scoring methods: JASMINE^8^ (using the likelihood or odds-ratio settings, Jasmine_LH and Jasmine_OR, respectively), UCell^13^, the original method by Tirosh et al.^17^ implemented in Seurat^11^ (Seurat), a similar method implemented in Scanpy^9^ (Scanpy), its alternative that used all genes as control genes (Seurat_AG), an alternative that used least variable genes as control genes (Seurat_LVG), and our proposed method Adjusted Neighbourhood Scoring (ANS). Like Seurat and Scanpy, Seurat_AG and Seurat_LVG first compute the average expression of the genes, sort them in increasing order, and divide them into 25 equally sized expression bins. While Seurat_AG selects all genes from an expression bin as control genes, Seurat_LVG chooses genes with the smallest dispersion. Conversely, ANS avoids binning and selects control genes in an average-expression neighborhood closest to the signature genes’ average expression values (Methods). We implemented all methods originally developed for the R language in Python (Figure 1a) and showed their identical behavior (Figure 1b, Methods). The Python package complementing this work provides access to all implemented methods.

**Figure 1.**
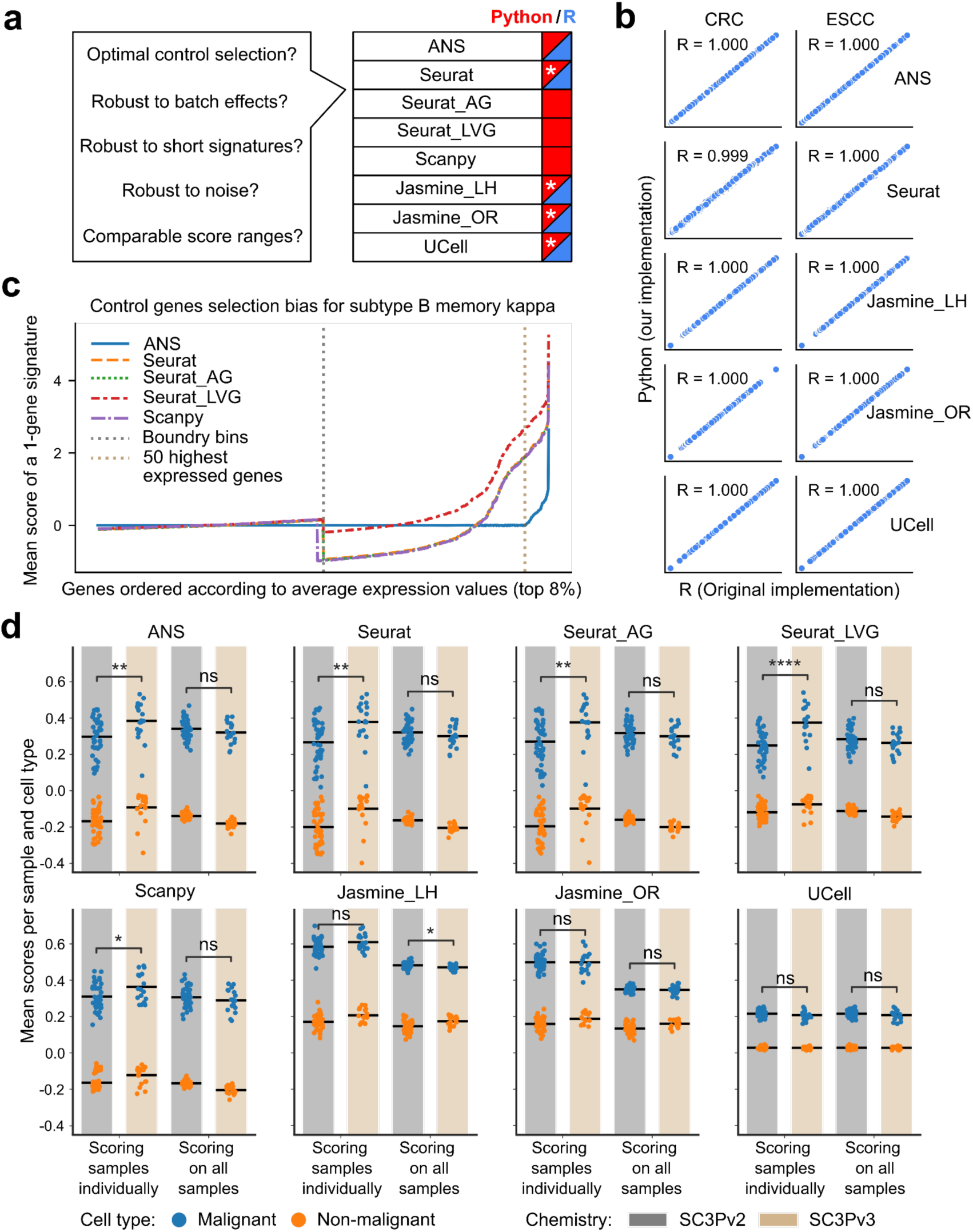
Performance and robustness analysis of gene signature scoring methods. **a,** Benchmark questions for the gene signature scoring methods and availability of the methods in Python (blue) and R (red). **b,** The scores obtained from R and Python implementations compared for two cancer datasets (CRC, ESCC). Discrepancies in Seurat scores between R (AddModuleScore of Seurat) and Python are attributed to randomization in the scoring method. **c,** Control gene selection bias of the scoring methods for the top 8% of highly expressed genes in B memory kappa cells within the PBMC dataset. The x-axis shows genes sorted by their average expression levels; the y-axis shows the mean score across all cells for a single-gene signature. Vertical dashed lines indicate the expression bin boundary and the top 50 highly expressed genes. The bias of a scoring method is indicated by how far the mean score of a gene deviates from zero. **d,** The influence of dataset composition and batch effect on scoring CRC cells using a 100-gene signature associated with malignant cells. Each dot represents the mean score for all cells within a sample, grouped by cell type (malignant in blue or non-malignant in orange), sequencing chemistry type (SC3Pv2 in grey or SC3Pv3 in beige), and scoring mode (scoring all the samples together or individually). The black horizontal bar represents the mean value of all dots within each group. P-value annotation: ns: p > 0.05, *: 0.01 < p <= 0.05, **: 1e-3 < p <= 0.01, ***: 1e-4 < p <= 1e-3, ****: p <= 1e-4.

We benchmarked the methods in six scenarios (Figure 1a, Table 1). In the first scenario, we showed the implication of control gene selection on gene signature scoring. In the second and third, we investigated the influence of the data composition and batch effects on scoring. In the fourth and fifth scenarios, we analyzed the impact of the variability in the gene signature length on the resulting scores. In the sixth scenario, we examined the scores produced by different methods when scoring multiple gene signatures and evaluated the performance of cell score-based classifiers to annotate cell types and cell states. The initial scenario involved a preprocessed dataset of peripheral blood mononuclear cells^20^ (PBMC); the next four scenarios were carried out on preprocessed datasets of colorectal carcinoma^21^ (CRC) and squamous cell carcinoma of the esophagus^1^ (ESCC); the sixth scenario used PBMC scRNA-seq data^20^ and scRNA-seq data from four cancer types: breast carcinoma^22^ (BRCA), high-grade serous ovarian cancer^23^ (HGSOC), cutaneous squamous cell carcinoma^24^(cSCC), and lung adenocarcinoma^25^ (LUAD) (Methods).

**Table 1.** Qualitative comparison of signature scoring methods based on their properties, influencing factors, and score comparability. ✓ means a scoring method fulfills property or resists an influencing factor. ✕ means the opposite. NA stands for a property or an influencing factor that cannot be evaluated for a scoring method. These results are discussed in detail in Results.

| Scoring method | Deterministic | Optimal control genes selection | Independence of total cell number and cell type proportions | Robust to batch effect when scoring all samples together | Robust when scoring for small signatures | Robust to unrelated genes in the signature | Provides comparable scores for multiple cell-type signatures |
| --- | --- | --- | --- | --- | --- | --- | --- |
| ANS: Adjusted Neighborhood Scoring | ✓ | ✓ | ✗ | ✓ | ✓ | ✓ | ✓ |
| Scanpy<sup>9</sup> : based on Tirosh *et al.*<sup>17</sup> | ✗ | ✗ | ✗ | ✓ | ✓ | ✓ | ✗ |
| Seurat<sup>11</sup>: based on Tirosh *et al.*<sup>17</sup> | ✗ | ✗ | ✗ | ✓ | ✓ | ✓ | ✗ |
| Seurat_AG: Seurat scoring with all genes as control | ✓ | ✗ | ✗ | ✓ | ✓ | ✓ | ✗ |
| Seurat_LVG: Seurat scoring with least variable genes as control | ✓ | ✗ | ✗ | ✓ | ✓ | ✗ | ✗ |
| Jasmine_LH: JASMINE with likelihood computation<sup>8</sup> | ✓ | NA | ✗ | ✓ | ✓ | ✓ | ✗ |
| Jasmine_OR: JASMINE with odds-ratio computation<sup>8</sup> | ✓ | NA | ✗ | ✓ | ✗ | ✓ | ✗ |
| UCell<sup>13</sup> | ✓ | NA | ✓ | ✓ | ✓ | ✓ | ✗ |

We also evaluated the time requirements for the eight methods as a function of the increasing number of cells and the increasing number of genes in the signature (Suppl. Figure S2). All methods exhibited linear scalability with dataset size. For short signatures (up to 10 genes), Seurat, Scanpy, and ANS performed the fastest (<10 sec for 150,000 cells), while Seurat and Scanpy remained the quickest for longer signatures (1,000 genes). In contrast, UCell and Jasmine took approximately seven times longer, while ANS took about three times longer in this case. Nevertheless, these methods consistently operated within seconds to a few minutes, even with large signatures and datasets.

### Control genes selection bias in gene signature scoring

All scoring methods based on Tirosh *et al.*^17^, *i.e.*, all methods except JASMINE and UCell, use control genes for gene signature scoring. Typically, genes whose average expressions are similar to the average expression of the signature gene are used as control genes. Consequently, these control genes exhibit a mean average expression, referred to as the mean control value, that is the closest approximation to the average expression of the signature gene itself. The selection of an optimal control set is crucial, as inappropriate selection can introduce a bias and lead to unreliable scores for all cells. We compared strategies for the control gene selection of the methods ANS, Seurat, Seurat_AG, and Seurat_LVG across diverse preprocessed subsets of cell subtypes within the PBMC dataset for the top expressed genes where the effects of the control gene selection were expected to be the most pronounced; we chose the 8% of genes with the highest average expression values corresponding to the last two bins of the Tirosh-based methods (Figure 1c, Suppl. Figure S3, Methods). We scored each gene in the last two expression bins (top ∼900 genes) for each scoring method. On homogeneous datasets, scores of single-gene signatures are expected to be distributed around zero. While we observed minimal score biases in the last but one expression bin, the last bin showed large biases in the score values for all published methods due to the significant variance of the genes’ average expressions. Whereas Seurat, Seurat_AG, and Scanpy first underestimated two-thirds of the gene scores and overestimated one third, Seurat_LVG overestimated almost all scores in the last expression bin (Figure 1c). All methods showed notable score biases for the top fifty expressed genes; these genes get typically excluded from gene signatures by ANS due to the impossibility of constructing a valid control gene set. Overall, ANS induced the smallest bias and thus used the optimal control gene sets.

### The influence of cell type proportions, batch effects, signature length, and inclusion of irrelevant genes on gene signature scoring

The mean control values and, thus, the choice of control genes depend on the dataset composition of the tissue or sample analyzed. Consequently, when utilizing methods that employ control genes, we anticipated greater variability in gene signature scoring values when scoring was conducted on a per-sample basis compared to a scenario when all cells from all samples were assessed collectively.

To highlight this effect, we calculated scores of 100-gene signatures associated with the malignant cell phenotype of CRC and ESCC (Methods). We compared the distributions of the mean scores per sample and cell type under two scenarios: when scoring was conducted on a per-sample basis and on all samples together (Figure 1d, Suppl. Figure S4). We observed a reduction of score variance for all scoring methods and both CRC and ESCC datasets, except for Seurat_LVG and UCell in the ESCC dataset, when scoring was performed on cells of all the samples together (Suppl. Figure S5), leading to more comparable score ranges over all samples. Additionally, we assessed how scoring on all samples together contributed to diminishing the batch effects for the Tirosh-based scoring methods. For this, we evaluated the differences in score distributions across two batches representing different sequencing chemistry, called SC3Pv2 and SC3Pv3 (Figure 1d). Batch effects significantly affected the score values of all Tirosh-based scoring methods when the scoring procedure was performed on a per-sample basis. In contrast, we observed decreased distribution shifts between chemistry-related batches when scoring on all samples. Therefore, our results suggest that scoring should be performed on all cells in the dataset simultaneously for all methods using control gene sets, including ANS.

Next, we assessed scoring methods’ robustness to changes in the signature length using CRC and ESCC datasets. Methods were compared based on the minimum number of signature genes needed to perfectly classify malignant and non-malignant cells (AUCROC, Methods). Most methods, including ANS, achieved perfect classification with only 12-15 top differentially expressed genes chosen on both datasets as genes with the highest log2FC and significant adjusted p-values (Suppl. Figure S4 and S6). The only outlier, Jasmine_OR, required ∼11 times more genes for an accurate cell type annotation in CRC. The small length of these effective signatures might be due to the relative ease of distinguishing malignant from non-malignant cells, *i.e.*, that the signatures contain genes with strong separating signals between the two populations. This can also be attributed to the signature selection process (Methods).

To test the methods’ robustness to the presence of irrelevant genes in a signature, we progressively replaced genes in a “pure” 100-gene malignant signature with random genes (Methods). All methods, including ANS and except Seurat_LVG, maintained high performance (AUCROC > 0.9) with up to 85% noise, requiring only ∼15 informative genes (Suppl. Figure S4 and S6). Seurat_LVG consistently underperformed, needing at least twice as many correct genes for comparable results in both datasets.

### Assessing score-range comparability for score-based cell type and state annotation

The most important application of gene signature scoring is to measure the activation of different transcriptional programs and use the scores to make statements about cells’ association with cell states or cell types. However, interpreting cell scores and assigning cells to states or types requires gene signatures that yield scores with high information quantity and comparable score ranges. To assess the information quantity and the comparability of score ranges produced by ANS and other scoring methods, we analyzed the accuracy of cell-type and cell-state annotation based on gene signature scores produced by different methods across four cancer datasets (BRCA, HGSOC, LUAD, and sCC) and four PBMC subsets^20^. We used cell-type or cell-state-specific non-overlapping gene signatures provided with the datasets (cancer) or selected gene signatures based on a provided list of differentially expressed genes per subtype (PBMC dataset, Methods)(Suppl. Table S1). The fairness of produced scores was estimated by evaluating the accuracy of an unsupervised cell annotation based on the argmax assignment of cells to states from signature scores. Additionally, we evaluated the information quantity of scores produced by each scoring method using a cross-validated supervised logistic regression predicting cell state based on signature scores (Methods).

We observed that score-based cell annotation—labeling cells with the cell type or state corresponding to the highest scoring signature—was accurate for distinguishing cell types (B cells, monocytes, and NK cells) in the PBMC dataset using scores produced by most methods (Figure 2a). All methods achieved high balanced accuracies ranging from 0.921 (Jasmine_OR) to 0.999 (Scanpy and UCell) and F1-scores from 0.958 (Seurat) to 0.999 (UCell), demonstrating robust performance in distinguishing these distinct cell types (Suppl. Table S2). Indeed, most scoring methods produced comparable score ranges, with the highest scores being observed for cells of matched cell type; JASMINE was an exception, showing overlapping ranges for B-cell and other signatures in B cells (Figure 2a).

**Figure 2.**
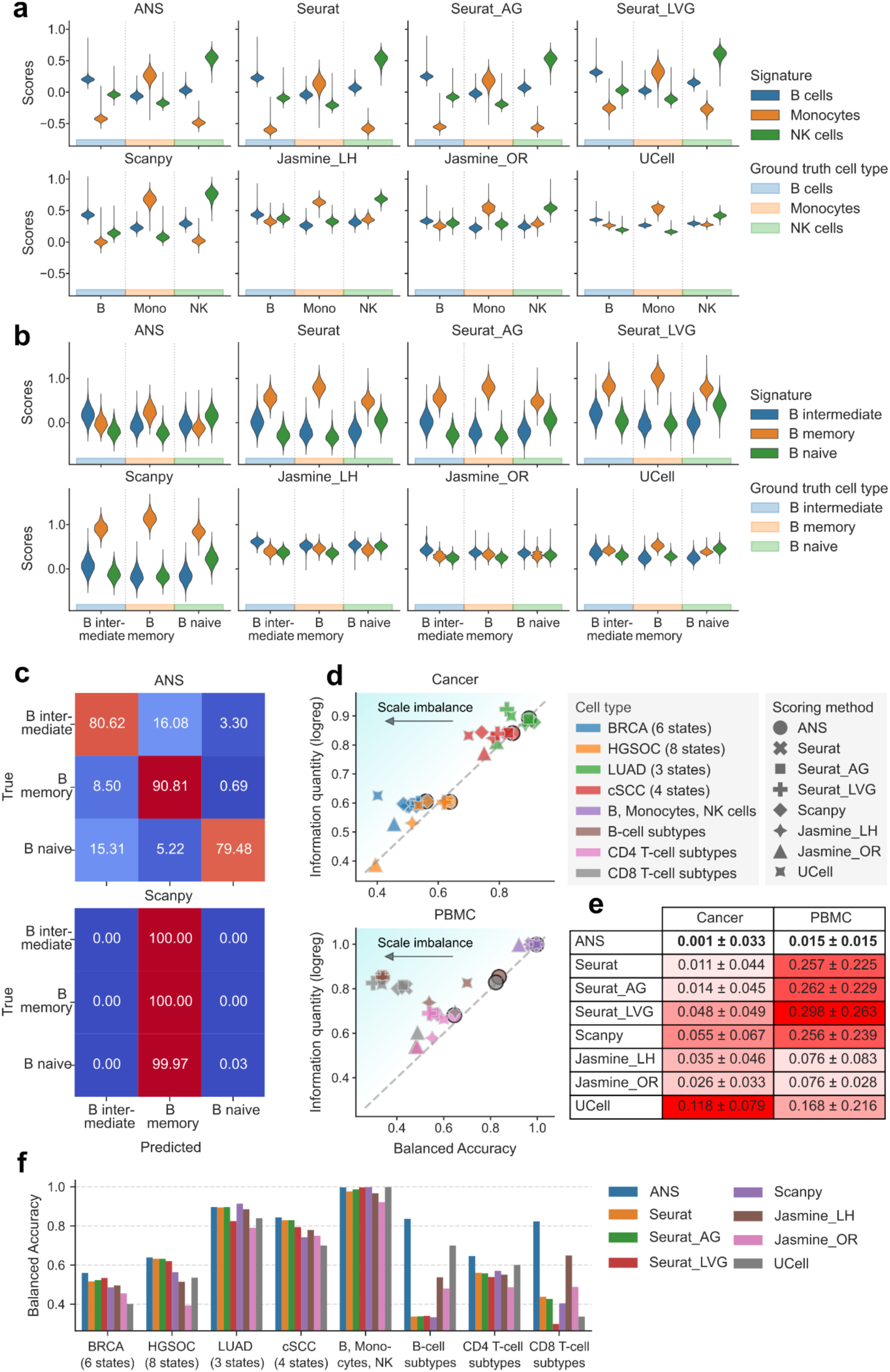
Comparative analysis of scoring methods for cell type and state annotation. Only non-overlapping cell type- or state-specific signatures were used. **a,** Score distributions for cell-type-specific signatures (B cells, monocytes, and NK cells) separated by true cell type annotations, calculated for each scoring method. **b**, Score distributions for B-cell subtype signatures separated by true cell subtypes, calculated by each scoring method. **c**, Row-normalized confusion matrix of B-cell subtype annotation based on the highest scores. **d**, Relationship between hard labeling performance and score information quantity in cancer vs. PBMC datasets. Scatterplots show balanced accuracy (x-axis) against score information quantity (y-axis) for various scoring method-dataset combinations. Balanced accuracy quantifies hard labeling performance, while score information quantity indicates the scores’ effectiveness in subtype classification. The diagonal line indicates perfect metric alignment, with vertical distances from this line representing scale imbalance. **e**, Quantitative analysis of scale imbalance across scoring methods and tissue types (cancer vs. PBMC). The mean and standard deviation of scale imbalance for each method are shown. Scale imbalance is the absolute difference between score information quantity and balanced accuracy in direct label assignment. The method with the lowest mean scale imbalance, indicating optimal consistency between information content and labeling accuracy, is highlighted in bold. **f**, Cell-state and -type annotation performance overview for all eight datasets and scoring methods.

However, when we applied the score-based annotation approach to assigning cells to cell states— naive, intermediate, and memory B-cells—we revealed limitations in using scores produced by all methods except ANS. Indeed, ANS was the only approach to produce informative and comparable score ranges for matched cell types that can be used for cell state classification based on the cell assignment to the state with the highest score (Figure 2b). Tirosh-based methods (Seurat, Seurat_AG, Seurat_LVG, and Scanpy) consistently scored highest for the B memory cell signature, regardless of the true cell type, resulting in poor labeling performance (Figure 2c, Suppl. Figure S7). UCell showed higher B memory cell scores than B-intermediate-cell scores when applied to B intermediate cells. Conversely, JASMINE scored highest for the B-intermediate-cell signature in B intermediate cells but failed to produce distinguishable score ranges for other cells.

To comprehensively evaluate this pattern, we expanded our analysis and assessed the performance of the score-based cell state classifier on four datasets of malignant cells with ground-truth assignments of cells to malignant cell states (BRCA, HGSOC, LUAD, and sCC) and two additional PBMC subsets comprising CD4 and CD8 cell state annotations. Across these six additional datasets, ANS maintained strong performance, providing unbiased gene signature scores that were useful for cell annotation, achieving the highest balanced accuracy in five cases (BRCA: 0.560, HGSOC: 0.638, cSCC: 0.843, CD4 T-cells: 0.645, CD8 T-cells: 0.823) and performing competitively in LUAD (0.897 vs. Scanpy’s 0.914)(Figure 2f, Suppl. Figure S7-S8, Suppl. Table S2).

We then compared the cell classification accuracy of the unsupervised score-based approach with a supervised approach of a trained score-based linear classifier. We refer to the cross-validated accuracy of the best linear classifier based on the gene signature scores as information quantity of the gene signatures (Methods). Information quantity close to 1 corresponded to the perfect supervised annotation of cell states or types based on gene signature scores. Methods with original score label assignment performance close to the information quantity were considered to produce fair gene signatures within comparable ranges, indicating a small score scale imbalance. In this experiment on eight datasets, ANS consistently ranked among the best-performing methods, showing the closest alignment of balance accuracy of the unsupervised score-based annotation with information quantity across (Figure 2d). Notably, ANS demonstrated substantially better performance for sCC, CD8, and B-cell subtype classification compared to other scoring methods.

Furthermore, across both cancer and PBMC datasets, ANS exhibited minimal scale imbalance measured as the difference between information quantity and balanced accuracy of score-based cell annotation, also with the smallest standard deviation, suggesting fairly comparable score ranges (Figure 2e, Suppl. Table S2). While these results are based on non-overlapping gene signatures to avoid bias from redundant genes, similar outcomes were observed for signatures that shared certain genes (Suppl. Figure S9-S10, Suppl. Table S3).

In conclusion, while all methods excel in discriminating distinct cell types based on gene signature scores, the task becomes significantly more challenging when discriminating signals from different cell states. ANS demonstrated the best performance and consistency across various cancer and healthy datasets by providing gene signature scores useful for unsupervised score-based cell annotation.

### Devising cancer cell-specific EMT signature: a case study for the ANS application

As we demonstrated the robustness and performance of the ANS method in the challenging scenario of gene signature scoring on related cell states, we chose to showcase the ability of ANS to help extract biologically relevant gene signatures in a similarly difficult setting. Specifically, we applied ANS to address the problem of the lack of malignant cell-specific EMT signature applicable across cancer types. The difficulty in constructing such a gene signature comes from the high similarity of transcriptional states of malignant EMT-exhibiting cells with malignant non-EMT cells, on the one hand, and with non-malignant mesenchymal cells, such as cancer-associated fibroblasts (CAFs), on the other hand. Building a cancer cell-specific signature for EMT would allow scoring bulk RNA-seq tumor datasets to quantify the degree of EMT transformation in human tumors while minimizing the bias induced by the presence of non-malignant cells of mesenchymal origin in the tumors.

Indeed, when we explored the behavior of previously published EMT signatures using the ANS scoring method on single cells from the ESCC^1^, lung adenocarcinoma^26^ (LUAD_Xing), CRC^21^ and breast carcinoma^22^ (BRCA) tumors, we observed that the pan-cancer EMT signatures, such as the Hallmark EMT^27^, the pEMT signature^28^, and five other EMT signatures^29–33^, resulted in extremely high scores in CAFs in addition to the mesenchymal-like (MES-like) malignant cells (Figure 3a and Suppl. Figure S8). This bias was likely due to the high expression in CAFs of such mesenchymal markers as *ACTA2*, *CDH2*, *FN1*, *SNAI2*, and *VIM* for the Hallmark EMT signature^34^, and *PDPN*, *TNC*, and *VIM* for the pEMT signature^35,36^. The other cancer EMT gene signatures produced high scores also for other cell types, *e.g*., the signatures of Foroutan *et al.*^32^, Gröger *et al.*^29^, Mak *et al.*^31^, and Tan *et al.*^30^ resulted in high scores for pericytes in CRC and ESCC, and the former also scored high in endothelial cells for all datasets (Suppl. Figure S8).

**Figure 3.**
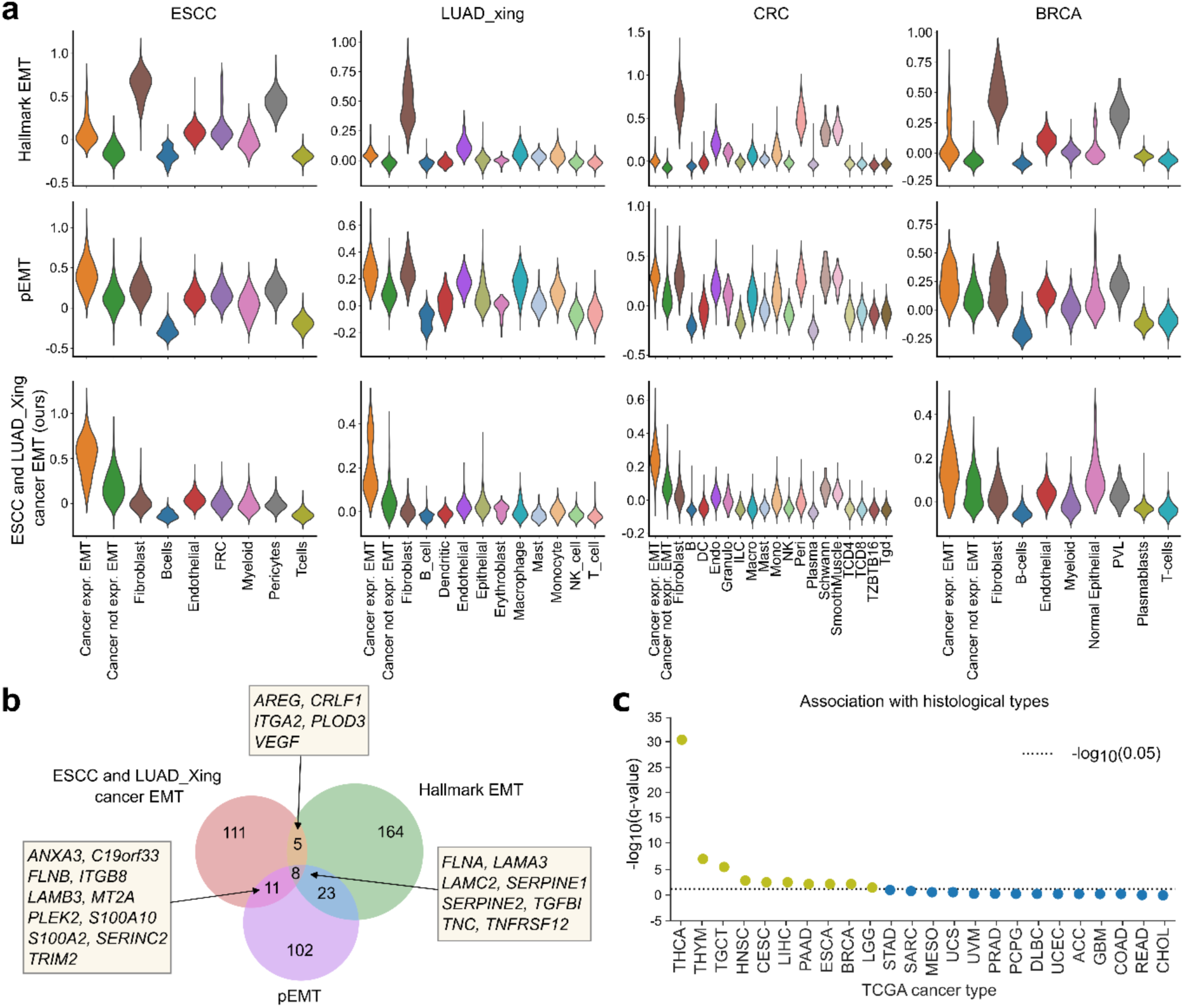
Application of ANS to devise EMT signature specific to malignant cells. **a**, Score distributions of the Hallmark EMT, the pEMT, and our proposed ESCC- and LUAD_Xing-specific cancer EMT gene signatures in different cell types of ESCC, LUAD_Xing, CRC, and BRCA. **b**, Venn diagram for the overlap in gene lists for the Hallmark EMT, pEMT signatures, and the signature we designed. **c**, Association of ESCC- and LUAD_Xing-specific cancer EMT signature scores and histological subtypes in TCGA (only cancer types with at least 1 histotype are included). Significant associations (qvalue < 0.05) are represented by dots above the dotted line. The FDR- adjusted p-values from the Kruskal-Wallis test were −log_10 transformed and sorted in decreasing value, with a higher y-value indicating higher significance.

To build a cancer cell-specific EMT signature, we leveraged two public scRNA-seq datasets, the ESCC and LUAD_Xing, for training and then validated the signature on the single cells of CRC and BRCA. To devise a cancer-specific EMT signature that scores high for MES-like malignant cells but low for non-malignant CAFs and malignant cells not undergoing EMT, we extracted genes with the abovementioned properties in ESCC and LUAD_Xing single-cell datasets (Methods). The devised cancer cell-specific EMT signature included many known EMT-related genes. We found 8 of the 135 signature genes intersecting with the Hallmark EMT and the pEMT signatures (Figure 3b). Moreover, 43 out of 135 genes appeared to be in at least one of the previously considered pan-cancer EMT signatures (Suppl. Table S4). By construction, the devised signature scored highly for the MES-like malignant cells in these two cancer types used for training (Figure 3a). We observed its strong discriminative power on the validation datasets CRC and BREAST (Figure 3a, Suppl. Table S5). The accuracy of our gene signature discriminating MES-like malignant cells versus all remaining cells measured with the area under the precision-recall curve (AUCPRC) on CRC and BRCA datasets was 0.701 and 0.511, respectively; it was slightly outperformed on CRC only by the gene signature proposed by Hollern *et al.*^33^ (AUCPRC 0.707) (Suppl. Table S5). However, this signature performed sub-optimally in our ESCC and LUAD_Xing training datasets (AUCPRC 0.028 versus 0.592 of our gene signature in ESCC, and 0.244 versus 0.513 in LUAD_Xing).

To investigate the clinical significance of the proposed cancer cell-specific EMT signature, we used bulk RNA-seq and clinical information from The Cancer Genome Atlas (TCGA) datasets. Specifically, we were interested in analyzing the link between the extent of cancer cells expressing EMT and the molecular subtypes of analyzed samples. Calculation of cancer cell-specific phenotypes using bulk RNA-seq data poses a significant advantage in cases when corresponding single-cell data is not available for a number of reasons. First, since EMT plays a crucial role as one of the main drivers of tumor plasticity and invasiveness, accurately measuring its activity in both TME and cancer cells is essential for designing successful targeted therapies. Second, an association between the cancer cell-specific EMT activity and histological subtypes may highlight the role of the cancer cell EMT phenotype in promoting, likely more invasive, molecular subtypes, and expand our understanding thereof. We selected 24 TCGA datasets for which more than one histological subtype was available, scored the cancer cell-specific EMT signature, and associated the scores with histological subtypes (independently in each cancer type). We found our signature to be significantly associated with the histological subtype of 10 cancer datasets (Figure 3c, Supp. Table S6). These results highlight the biological relevance of the proposed signature and its potential utility in further exploration of cancer molecular subtypes.

This example demonstrated how gene signature scoring on single-cell data could be applied to design meaningful gene signatures and annotate cells according to their transcriptional states for further biological exploration.

## Discussion

In the present work, we developed a deterministic and robust gene signature scoring method, Adjusted Neighborhood Scoring (ANS), which improves the approach introduced by Tirosh *et al.*^17^. In contrast to the Tirosh approach, ANS avoids binning genes by mean gene expression values and, by construction, chooses the optimal control gene sets to normalize gene expression values for technical effects. We demonstrated that the issue of sub-optimal stability of ANS and all methods based on the Tirosh approach, such as Scanpy and Seurat, when scoring gene signatures on samples with variable proportions of different cell types, can be solved by scoring all cells simultaneously using all samples in the dataset.

We compared ANS with other scoring methods developed for single-cell data to demonstrate the robustness and stability of the proposed scoring procedure. All scoring methods were shown to be stable upon scoring small signatures, and to be robust against noise, allowing up to 85% of noise in the signature if the remaining genes belong to the most differentially expressed gene discriminating the cell types tested. ANS was the top-performing approach in terms of providing high information quantity and comparable ranges of calculated scores for matched gene signatures and cell types. The remaining Tirosh-based scoring approaches showed high information quantity in their scores. However, they suffered from large scale imbalances, possibly influenced by the biased selection of the control sets and lack of score normalization. We also showed that scale imbalance could be avoided in Tirosh-based scoring methods through unsupervised clustering, converting signature scores to cluster assignments correlating with the matched cell state. Nonetheless, it is worth noting that the underlying assumption of exclusive state activation within individual cells could be limiting in certain situations. Furthermore, we have included all scRNA-seq signature scoring techniques within a signature scoring package, enabling the utilization of methods originally implemented in R in Python.

It is important to emphasize that gene signature scoring is only as reliable as the provided signatures. To generate confident and biologically meaningful scores, the signature must accurately capture the biological phenotype of interest. This is especially important when using signature scores for cell labeling, as an unreliable signature can result in misclassification. Poor gene signatures inevitably lead to poor labeling performance, as observed in BRCA, HGSOC, and CD4 signatures (Figure 2d). When using related gene signatures, scores may appear similar, which makes the selection of optimal control genes a critical step for distinguishing between closely related states and ensuring accurate results. ANS fulfills this need with a reliable control gene selection strategy, ensuring greater accuracy compared to other tested methods.

We showcased the use of the ANS scoring method to derive a gene signature specific to MES-like malignant cells that we validated in CRC and BRCA. Building a cancer cell-specific signature for EMT could allow scoring bulk RNA-seq tumor datasets to quantify the degree of EMT transformation in human tumors while minimizing the bias induced by the presence of non-malignant cells of mesenchymal origin in the tumors. Our ESCC- and LUAD_Xing-specific cancer EMT signature outperformed others in distinguishing MES-like malignant cells from CAFs across all datasets. When comparing MES-like malignant cells with other malignant cells, our signature ranked among the top four in terms of AUCPR for ESCC, LUAD_Xing, and CRC but placed sixth for BRCA (Suppl. Table S4). Notably, the Hallmark EMT^27^ signature displayed the highest discriminatory power in this context. The difficulty in discrimination may arise from the signature’s reliance on epithelial markers, lacking strong mesenchymal-related markers, which results in closer scores between malignant epithelial cells. Our cancer MES-like signature included a transcriptional factor, which so far has not been considered in any of the used pan-cancer EMT signatures: *PITX1*. *PITX1* encodes a member of the RIEG/PITX homeobox family and plays a crucial role in organ development and left-right asymmetry^37,38^. Previous research on *PITX1* in cancer has revealed cancer-type-specific effects. While upregulation of *PITX1* has been shown to be associated with tumor progression in lung adenocarcinoma^39^, kidney renal clear cell carcinoma^40^, breast^41^, epithelial ovarian^42^, and prostate cancer^43^, interestingly, in other cancer types, such as melanoma^44^, ESCC ^45^, osteosarcoma^46^ and head and neck squamous cell carcinoma^47^, down-regulation of *PITX1* has been found to enhance tumor progression. Other EMT-related genes exclusively present in our signature included two long-non-coding RNA genes: *ABHD11-AS1* and *BCYRN1*. *ABHD11-AS1* has previously been suggested as a prognostic biomarker for pancreatic cancer^48^, while *BCYRN1* was shown to promote cell migration and invasion in lung and colorectal cancers^49–52^. Additionally, the protein-coding genes *FAM83A*, *ITGA3*, *ITGB4*, *L1CAM*, *MUC16*, and *SAA1* have been reported to promote cancer progression in CRC^53,54^, lung cancer^55,56^, head-neck squamous cell carcinoma^57,58^, breast^59^, ovarian^60,61^ and endometrial cancer^62^. Lastly, our analysis has revealed that the identified cancer cell-specific EMT signature corresponds to histological subtypes in several TCGA datasets, highlighting the potential role of EMT-expressing cancer cells in their molecular characteristics. This finding highlights the biological significance of the proposed signature; however, future work is needed to verify its clinical applicability.

We acknowledge the limitations inherent in our study. Due to the exclusive assignment of each cell to a specific cell type and the difficulty in capturing cells transitioning between states—such as from naive B cells to memory B cells—our benchmarking did not include evaluating performance for such transitions. An alternative approach for such scenarios could involve associating scores with cell trajectories. However, our primary objective in this work was to assess the scoring specifically for cell annotation. Also, when applying our method to establish the cancer-specific EMT signature in the three datasets, LUAD_Xing, ESCC, and CRC, we relied on the selection of the cancer cells expressing previously published EMT-related genes. In contrast, in the basal-like breast cancer dataset, we relied on the original annotation of cell states and used the GM3 state strongly associated with EMT to label the EMT-presenting malignant cells. The selection procedure might have biased the downstream analysis.

In summary, this work presents an improved version of the scoring method proposed by Tirosh^17^, called Adjusted Neighborhood Scoring (ANS). ANS is a deterministic and robust scoring method that outputs comparable scores for multiple gene expression signatures, which can be used for cell type and cell state annotation in an unsupervised way. The ANS method is implemented in a user-friendly way to be integrated into R and Python single-cell analysis pipelines. By employing this technique, one can confidently evaluate cellular programs and develop highly specific cell state signatures, as exemplified by the proposed cancer EMT signature.

## Methods

### Dataset and Preprocessing

We used eight different single-cell RNA sequencing (scRNA-seq) datasets, comprising seven cancer datasets and one peripheral blood mononuclear cell (PBMC) dataset^20^. For control genes selection and basic robustness experiments, we utilized a squamous cell carcinoma of the esophagus^1^ (ESCC, 54 samples) and a colorectal carcinoma^21^ (CRC, 60 samples) dataset. To assess signature score range comparability, we considered malignant cells from four cancer datasets: breast carcinoma^22^ (BRCA, 20 samples), high-grade serous ovarian cancer^23^ (HGSOC, 134 samples), cutaneous squamous cell carcinoma^24^(cSCC, 7 samples), and lung adenocarcinoma^25^ (LUAD, 9 samples), as well as the PBMC dataset consisting of 24 samples collected at three different time points from eight patients. The case study included the ESCC and CRC datasets, along with another lung adenocarcinoma dataset^26^ (LUAD_Xing, 19 samples including 12 from subsolid-nodules and 7 from primary lung adenocarcinoma) and the 7 basal-like samples from BRCA. Cell type distributions for each sample used in the case study are presented in Suppl. Figure S12.

Preprocessing scRNA-seq data is crucial to address inherent biases and enable downstream analysis. Cancer datasets and the PBMC dataset were preprocessed differently. For the cancer datasets, except HGSOC,we used the preprocessing module of `CanSig`^63^ with the default parameter as the first preprocessing step (https://github.com/BoevaLab/CanSig). In brief, we applied standard quality control metrics and removed potential misannotated cells using CNV inference; the preprocessing steps are extensively described in the original paper. Additionally, we filtered genes based on a minimum expression in 1% of cells, and normalized cell reads using a shift logarithm log_2_(x+1) with the mean total raw count per cell as the target sum, as described and implemented by Ahlmann-Eltze and Huber^64^. For HGSOC, due to its size, we directly applied the 1% gene filtering and log-normalization on the cancer cells from the GEO data (GSE180661), skipping the CanSig preprocessing step. We demonstrated that these preprocessing choices are robust, as replacing the CanSig preprocessing pipeline with alternative approaches (e.g., Curated Cancer Cell Atlas preprocessing, https://www.weizmann.ac.il/sites/3CA/) or omitting it did not significantly affect the relative performance of the scoring methods (Suppl. Figure S13-S15). Regarding the PBMC dataset, we created several sub-datasets based on cell type and state annotations. The first sub-dataset consisted of B-cells, monocytes, and natural killer cells, while the second subdataset contained B-cell states (B-naive, B-intermediate, and B-memory cells). The third and fourth sub-datasets consist of CD4 T-cell states (CTL, naive, proliferating, TCM, TEM, and Treg) and CD8 T-cell states (naive, proliferating, TCM, TEM).

All sub-datasets were preprocessed as follows: Initially, we filtered low-quality reads according to the guidelines provided in chapter 6.3 of the book “Single-cell Best Practices” by Heumos *et al.*^65^. A threshold of 5 median absolute deviations (MADs) was selected for filtering covariates, including total counts, number of genes with positive counts in a cell, and the percentage of mitochondrial counts. Similar to the cancer datasets, we filtered genes based on a minimum expression in 1% of cells and normalized the cell reads using the shift logarithm with the mean total raw count per cell as the target sum.

### Gene signature selection for malignant cells in benchmark

To select gene signatures for malignant cells in CRC and ESCC, we adopted the following approach. To address data imbalance caused by varying sample sizes (Suppl. Figure S11), we bulkified the datasets per sample using the method `get_pseudobulk` from the Python package decoupleR^66^ with the parameters `mode=sum`, `min_cells=10`, and `min_counts=1000`. Bulkifikation of the samples prevented the dominance of genes expressed in the majority sample and minimized batch effects. As described in the “Pseudo-bulk functional analysis” tutorial of `decoupleR`^66^ (version 1.4.0), we further filtered genes by expression using the default parameters. We then applied the `PyDESeq2`^67^ (version 0.3.3) workflow (Python version of `DESeq2`^68^), a differential gene expression analysis (DGEX) tool, to the bulkified samples. Using `DESeq2`, we identified genes with significant differential expression (log_2_FC > 2 and adjusted p-value < 0.01) between malignant and non-malignant cells. This process helped us pinpoint gene signatures strongly associated with the malignant cell phenotype in CRC, and ESCC, which were used during the benchmark.

### Control genes selection in Tirosh-based gene signature scoring and ANS

The methods implemented in the scRNA-seq analysis packages `Scanpy`^9^ (in Python) and `Seurat`^11^ (in R) were built upon the procedure first described by Tirosh *et al.*^17^ Here, we recall this procedure in a formal manner. Let 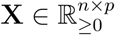 be the preprocessed expression matrix for *n* cells and *p* genes, generally representing log-transformed normalized read counts. Let *S* = {*s*_1_, …, *s_m_*} be a given gene signature, *i.e*., a set of *m* genes. Let 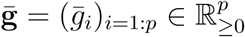 be the average expression vector for each gene *i* over all cells ordered by increasing average expression (i.e., *ḡ_i_* ≤ *ḡ_i_*+1 for any *i*).

In Tirosh-based gene signature scoring methods, ḡ is split into 25 equally sized bins, called expression bins. For each signature gene *s*, we sample *c* (control size) genes from *s*’s expression bin. Let *C_s_* be the set of control genes for the signature gene *s*. The score for a cell *j* is then computed as follows:

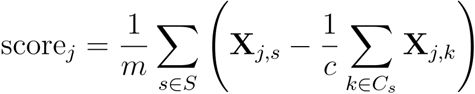

In this work, we kept the main idea of the scoring approach proposed by Tirosh *et al*. but suggested and validated three alternatives to select control genes.

When implementing the first two alternatives, we preserved the gene binning approach. In the “all genes as control genes” alternative (Seurat_AG), we selected all non-signature genes from an expression bin as control genes. Note that Seurat_AG was not using the control genes size parameter *c*. In the Least Variable control Genes alternative (Seurat_LVG), we first computed the least variable genes for each expression bin. The least variable genes were calculated with Scanpy’s `highly_variable_genes`^69^ method (flavor ‘seurat’) and selected by taking the *c* genes with the smallest dispersion.

In the third alternative, called Adjusted Neighborhood scoring (ANS), the control genes for a signature gene are selected based on the average expression neighborhood. Let *ḡ* be equal to *ḡ*, but excludes the signature genes *S*. Let 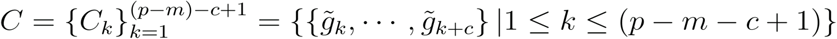 be the set of windows of size *c* (control size) of *ḡ*. For each signature gene *s*, we computed the control set as follows:

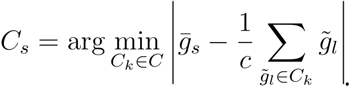

In other words, for each signature gene, we selected *c* control genes around this gene whose average mean expression closely matched the mean expression of the signature gene. Of note, ANS excluded signature genes within the 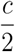 genes with the highest average expression to avoid invalid control gene selection.

### Evaluation of scoring methods for score-based cell type and cell state annotation

We performed unsupervised cell type/state annotation using gene signature scoring in all experiments. For a dataset with *n* signatures associated with *n* cell types/states, we assigned cell identity based on the highest scoring signature (argmax). Performance was assessed using `scikit- learn`^70^ (version 1.2.0.) metrics: AUCROC for balanced datasets, AUCPRC for unbalanced datasets, along with balanced accuracy and weighted F1-score.

### Scoring Methods in R and Python

We implemented the R methods UCell (https://github.com/carmonalab/UCell) and JASMINE (https://github.com/NNoureen/JASMINE), as well as the original method proposed by Tirosh et al.^17^, in Python. Additionally, we provided the ANS method in both Python and R. We showed that the scoring methods yield identical scores for both Python and R. We selected a 100-gene signature for malignant cells based on the genes with the lowest adjusted p-value and the highest log_2_FC. We then scored this signature on preprocessed datasets of two cancer types: CRC and ESCC. To ensure consistency, we converted these datasets into `SingleCellExperiments`^71^ and employed the original R versions of the scoring methods. Notably, the R implementation of the Seurat method based on the original version of Trisoh *et al.*, known as `AddModuleScore`^72^, is included in the scRNA-seq package `Seurat`^20^. To evaluate the concordance between the R and Python scores, we computed the Pearson correlation coefficient using the `scipy` package^73^ (version 1.9.3) in Python (Figure 1b). The implementations can be found in our software package (https://github.com/lciernik/ANS_signature_scoring).

### Benchmark experiments

In the first part of the study, we compared the properties and resistance to influencing factors of eight gene signature scoring methods: JASMINE^8^ using the likelihood or odds-ratio (Jasmine_LH and Jasmine_OR), UCell^13^, the original method by Tirosh et al.^17^ implemented in Seurat^11,12^ (Seurat), the method implemented in Scanpy^9^ (Scanpy), Seurat using all genes as control genes (Seurat_AG), Seurat using the least variable genes as control genes (Seurat_LVG), and our proposed method Adjusted Neighbourhood Scoring (ANS). Unless stated differently, we chose signatures of desired lengths associated with the malignant cell phenotype for the cancer datasets CRC and ESCC. This selection was based on identifying genes with the smallest adjusted p-value and the largest log_2_FC.

#### Control gene selection bias on Tirosh-based gene signature scoring

The Tirosh-based scoring methods rely on the selection of control genes. However, Seurat, Seurat_AG, Seurat_LVG, and Scanpy can introduce biases in scoring when there is substantial variability in gene expression averages within expression bins, which is especially visible in the top 8% of the highest expressed genes (Suppl. Figure S1). The ANS method was motivated to reduce the issue of bias induced by control genes. We compared the strategies for the control gene selection of the methods ANS, Seurat, Seurat_AG, and Seurat_LVG across the four preprocessed single-cell-type PBMC sub-datasets: B memory kappa, CD8 TM2, CD14 Mono, and NK 3 (Dataset preprocessing in Methods). For each dataset, we scored each gene in the top 8% of the highest expressed genes and, thus, converted the expression matrix to a score matrix. On homogeneous datasets, scores of a gene are expected to be distributed around zero. We visualized the mean scores and standard variation for each gene in increasing average expression order with the `lineplot` function of the plotting package `searborn`^74^ (Figure 1c, Suppl. Figure S3).

#### The influence of the cell type proportions and batch effects on gene signature scoring

The two following experiments explored the influence of dataset composition on gene signature scoring. The first experiment investigated the influence of different cell type proportions. The second experiment tested the influence of batch effects on scoring methods. Both experiments used the preprocessed CRC dataset and a 100-gene signature for malignant cells with the smallest adjusted p-values and the highest log_2_FCs. The CRC dataset contains samples sequenced with two different sequencing chemistries. We used the different sequencing chemistries as surrogates for batch effects. We scored each sample individually and the entire dataset. For each scoring method, we averaged the cell scores for each sample, cell type (malignant or non-malignant), scoring mode (scoring samples individually or all together), and sequencing chemistry (SC3Pv2 or SC3Pv3). We compared the variance between the sample score averages for each scoring method and scoring mode in the first experiment and observed reduced or equal variance when scoring on the entire dataset. In order to evaluate the impact of batch effects, we utilized Mann-Whitney U testing to compare the sample score averages of malignant cells between the two sequencing chemistries for each scoring method and scoring mode. We used the Python package `statannotations`^75^ (version 0.4.4.) to compute the Mann-Whitney U tests and add statistical annotations in Figure 1d. Scoring samples individually showed significant differences in score ranges between chemistries. We performed identical experiments on the preprocessed ESCC dataset, excluding the batch effect component because of using uniform sequencing chemistries for all samples in those datasets (Suppl. Figure S4).

#### The influence of signature length on gene signature scoring

The next experiment aimed to assess the robustness of scoring methods to small signatures. Specifically, we sought to determine the minimum number of genes necessary to achieve perfect discrimination between malignant and non-malignant genes using scores, thereby enabling binary classification. For each scoring method, we followed a systematic approach. We began by selecting a base signature for the malignant phenotype. Subsequently, we removed genes for which a valid control set could not be constructed, specifically those belonging to the 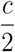 smallest or highest expressed genes. The remaining genes were then sorted based on ascending adjusted p-value and descending log_2_FC. We iterated over the signature to ascertain the number of genes required, progressively expanding the range of elements considered from the initial subset containing only the first element until the final iteration encompassing the entire list. Utilizing the methodology outlined in the Method section *Evaluation of scoring methods*, we computed the area under the receiver operating characteristic curve (AUCROC) for each signature length, using both the scores and malignancy annotations. We ceased computing as soon as an AUCROC value of 1 was achieved. We conducted the analysis for the preprocessed CRC and ESCC datasets (Suppl. Figure S4 & S6).

#### Robustness to noise in gene expression signature

To compare the robustness of the gene signature scoring methods to noise, we utilized a 100-gene base signature, which achieved an AUCROC of 1 for all scoring methods and iteratively replaced genes by noise. We selected noise genes based on their adjusted p-value > 0.01 and their |log2FC| <= 0.5 during DGEX, indicating that these genes lacked statistical significance in distinguishing malignant and non-malignant cells. During each simulation run, we iteratively and randomly substituted the signature genes with noise genes, starting from the pure signature and progressing until the entire signature consisted of noise genes. We conducted 20 simulation runs for each gene signature scoring method to ensure comprehensive results. The experiment was performed on preprocessed CRC and ESCC datasets (Suppl. Figure S4 & S6).

#### Comparability of signature score ranges for cell type and state annotation

To assess the information quantity and ranges of scores when scoring for multiple signatures, we considered unsupervised and score-based cell type and state annotation of four cancer datasets (BRCA, sCC, HGSOC, LUAD) and four PBMC dataset subsets. As previously mentioned, only malignant cells from cancer datasets have been used for the annotation task. We used cancer cell state signatures published with the datasets. For the PBMC datasets, we constructed cell type-/ state-specific signatures using differential gene expression data accompanying the scRNA-seq dataset (https://atlas.fredhutch.org/nygc/multimodal-pbmc/). The PBMC dataset annotates cells at three levels of granularity: level one for broad cell types, level two for intermediate subtypes, and level three for the most detailed subtypes. For each selected cell type in our PBMC subset datasets, we built signatures by identifying all corresponding level-3 subtypes and selecting differentially expressed genes (DEGs) with log2 fold change > 0.05 and p-value < 0.01. We created the final signature by combining DEGs from these level-3 subtypes through set union operations. When comparing overlapping versus non-overlapping signatures, we additionally created a version where genes appearing in multiple cell type signatures were removed, as these shared genes could bias scoring across cells.

For each dataset, we calculated scores for every cell using all signatures and scoring methods, each signature being associated with a distinct cell type/state (score distributions in Suppl. Figure S8). We assigned each cell to the state associated with that cell’s highest-scoring signature (argmax), enabling unsupervised annotation.

To evaluate the information content within the scores, we employed scikit-learn’s (v1.2.0) cross-validated supervised logistic regression (LogisticRegressionCV with C=None and stratified k-fold cross-validation where k=10, random_state=42) to assess the predictive power of the scores. We used balanced accuracy and weighted F1-score metrics to evaluate both supervised and unsupervised approaches. We defined scale imbalance as the difference between the cross-validated supervised classification performance and unsupervised score-based annotation performance. Results for non-overlapping and overlapping signatures are presented in Suppl. Tables S2 and S3, respectively.

#### Case study: selection of cancer EMT cells and cancer-specific EMT signature establishment

For the entire case study, we used ANS for evaluation with control sets of size c=100. The generation of the cancer-specific EMT signature consisted of two main steps. In the first step, we classified the malignant cells as EMT-expressing or non-expressing. Based on this classification, we explored the genetic differences between these two groups and between EMT cancer cells and cancer-associated fibroblasts.

#### Classification of cancer EMT cells

As a first step in the signature establishment, we selected cancer cells expressing EMT, called cancer EMT cells. We computed scores for ESCC with the dataset-specific mesenchymal signature^1^. We scored the entire dataset and the subset of malignant cells. Cancer cells being within the 10% highest scoring cells on the entire dataset and in the subset were classified as cancer EMT cells. The 3’860 cancer EMT cells out of 41’399 cancer cells (9.32%) and 178’109 total cells (2.17%), stem from 53 out of 54 samples. For LUAD_Xing and CRC, no dataset-specific mesenchymal signatures were reported. Therefore, we selected the cancer cells expressing EMT as follows. We scored the datasets for seven pan-cancer EMT signatures: Hallmark EMT signature ^27^, pEMT gene module ^28^, and five other EMT signatures ^29–33^. The pan-cancer EMT signatures were found on EMTome ^76^. For each pan-cancer signature, we ranked the cancer cells by ANS scores and computed for each cell the median rank. Finally, 10% of cancer cells with the smallest median signature ranks were classified as cancer EMT cells. We selected the median to avoid outlier ranks of the pan-cancer signature. For BRCA, we used the program labels provided by the original paper^22^ and excluded cells falling in the “grey zone” of the EMT program expression (“GM3” in the original paper). We considered only samples assigned to the “Basal” subtype and defined the “grey zone” to be cells not labeled with “GM3”, but having a signature score for GM3 above 0.15 (scoring was applied on all samples similar in the original paper). In the LUAD_Xing dataset, 822 out of 8’205 cancer (10.02%) and 55’659 total (1.48%) cells were classified as EMT. All 19 samples contained cancer EMT cells. In the CRC dataset, 4’328 out of 43’252 cancer (10.01%) and 140’090 total (3.09%) cells were classified as cancer EMT cells distributed over all 60 samples. In the BRCA dataset, 2031 out of 5115 cancer (39,71%) and 27458 total (7,40%) cells were classified as cancer EMT cells distributed over all 7 samples. Dataset compositions are shown in Suppl. Figure S8.

#### Gene signature refinement with scores of single signature genes

Gene signature refinement aims to remove genes from a signature to achieve a higher discriminating power of the signature scores. We mainly used gene signature refinement in the case study, however, it could be applied to any signature. During the refinement of a signature associated with a subset of cells, gene signature scoring is applied to each signature gene individually. A signature gene remained in the signature only if the scores for the desired subset of cells were significantly higher than those for another subset of interest. To test for distributional shifts, we used the Mann-Whitney U (`scipy`, version 1.9.3) with Benjamini-Hochberg correction for p-values (`statsmodels`, version 0.13.5)) with `alpha` 1e-5. The process required available cell annotations for the desired subsets of cells.

#### Establishment of a cancer-specific EMT signature in ESCC and LUAD_Xing

The signature establishment consisted of finding differentially expressed genes between cancer cells expressing EMT and CAFs as well as cancer cells not expressing EMT in ESCC and LUAD_Xing. The cancer EMT cell classification for all datasets is provided in the supplementary information. We first started by finding a cancer-specific signature for ESCC, then found one for LUAD_Xing. Finally, we unioned the ESCC and LUAD_Xing-specific cancer EMT signature and refined it again on ESCC. For differential gene expression (DGEX) analysis between two cell groups of interest, we used Wilcoxon rank-sum (Mann-Whitney U) tests ^77^ with tie correction and Benjamini-Hochberg correction implemented in Scanpy’s `rank_genes_groups`.

##### Cancer-specific EMT signature for ESCC

First, we created two subsets of the ESCC dataset: cancer EMT cells and CAFs and all cancer cells. Let *A* be the set of DGEX genes (*logFC* ≥ 2 and an adjusted p-value < 0.001) between cancer EMT cells and CAFs. Let *B* be the set of DGEX genes (*logFC* ≥ 1.5 and an adjusted p-value < 0.001) between cancer EMT cells and cancer cells not expressing EMT. We joined sets *A* and *B*, called gene set *C*, and removed all genes from *C* that are DGEX (*logFC* ≥ 2 and an adjusted p-value < 0.001) in T-and Myeloid cells versus cancer EMT cells. The reduced gene set *C*, called gene set *D*, scored high for many cancer cells and thus required more refinement to distinguish the cancer EMT cells from the other cancer cells. Let the set *X* be all cancer cells that scored above 0.2 for the gene set *D*. We selected all DGEX genes (*logFC* ≥ 1.5 and an adjusted p-value < 0.001), called gene set *E*, in *X* between cancer EMT cells and cancer cells not expressing EMT. We scored each gene in *E* − *D* (n=451) independently and kept genes with significantly higher scores for cancer EMT cells than for CAFs, cancer cells not expressing EMT, and the rest of the cells. We unioned the gene sets *D* and *E*, resulting in our ESCC-specific cancer EMT signature (signature refinement). The detailed procedure can be found in the Jupyter Notebook `find_cancer_emt_signature_ESCC.ipynb` in the Supplementary material GitHub repository.

##### Cancer-specific EMT signature for LUAD

Xing: For the LUAD_Xing-specific cancer EMT signature, we followed a procedure similar to that for ESCC. First we created two subsets of the LUAD_Xing dataset: cancer EMT cells and CAFs and all cancer cells. Let *A* be the set of DGEX genes (*logFC* ≥ 2 and an adjusted p-value < 0.001) between cancer EMT cells and CAFs. Let *B* be the set of DGEX genes (*logFC* ≥ 1 and an adjusted p-value < 0.001) between cancer EMT cells and cancer cells not expressing EMT. We unioned gene sets *A* and *B*, called gene set *C*. For each cell type *X* not equal to cancer EMT cells, we selected DGEX genes (*logFC* ≥ 1 and an adjusted p-value < 0.001) of *X* versus cancer EMT cells and removed them from gene set *C*. The resulting gene set corresponded to our LUAD_Xing-specific cancer EMT signature. The detailed procedure can be found in the Jupyter Notebook `find_cancer_emt_signature_LUAD.ipynb` in the supplementary material GitHub repository.

##### Joining cancer-specific EMT signatures for ESCC and LUAD_Xing and refinement on ESCC

To obtain a signature that can distinguish between cancer EMT cells and CAFs, and cancer EMT cells and cancer cells that do not express EMT, we unioned the ESCC- and LUAD_Xing-specific cancer EMT signatures. We scored for each gene in the unioned signature in ESCC (n=267) and only kept a gene if it had significantly higher scores for cancer EMT cells than for CAFs, cancer cells not expressing EMT, and the rest of the cells (signature refinement). We excluded the insignificant genes and incorporated the LUAD_Xing-specific cancer EMT signature genes absent in the preprocessed ESCC dataset and the ESCC-specific cancer EMT signature genes absent in the preprocessed LUAD_Xing dataset. Suppl. Table S4 shows the 135 ESCC- and LUAD_Xing-specific cancer EMT signature genes. The detailed procedure of signature union and refinement can be found in the Jupyter Notebook `union_ESCC_and_LUAD_specific_EMT_signature_and_refine_on_ESCC.ipynb` in the supplementary material github repository.

#### Validation of cancer-specific EMT signature

We applied the ESCC and LUAD_Xing-specific cancer EMT signature on the CRC and BRCA datasets for validation. For all datasets, we computed the AUCPRC (*Evaluation of scoring methods* in Methods) for the signature scores between cancer EMT cells and CAFS, cancer cells not expressing EMT, and all other cells (rest) (Figure 3a).

#### Correlation of cancer-specific EMT signature to histological types

To analyze the link between de novo EMT signature and histological subtype of TCGA datasets, we downloaded pan-cancer normalized RSEM (RNA-Seq by Expectation Maximization) gene expression data from the GDC portal (https://portal.gdc.cancer.gov/). Clinical information on the histological subtypes was retrieved from Liu et al.^78^. The EMT signature was scored in each corresponding cancer type for which more than one histological subtype was available. The signature was scored using the *ssgsea* method from the `GSVA` R package^7^. The association between EMT scores and histological subtypes was evaluated using the Kruskal-Wallis test with FDR correction. The analysis was performed using R Statistical Software (v4.1.2; R Core Team 2021).

### Statistical information

The statistical tools, methods, and threshold for each analysis are explicitly described with the results or detailed in the figure legends or Materials and Methods.

## Data availability

The preprocessed datasets for CRC, ESCC, LUAD_Xing, BRCA, sCC, HGSOC, LUAD, and PBMC and the used signatures can be downloaded: https://drive.google.com/drive/folders/10L2gqapJbyOn_MbrZRHQG--n0Xj7wIyg?usp=sharing. The raw transcript count matrices can be found at GEO: GSE178341 for CRC, GSE176078 for breast cancer, and GSE160269 for ESCC. The raw transcripts for LUAD_Xing can be found with the accession number HRA000154 at http://bigd.big.ac.cn/gsa-human. The raw transcripts for the remaining datasets can be found at GEO: sCC (GSE144240, GSE144236), HGSOC (GSE180661), LUAD (GSE131907). The raw PBMC dataset can be downloaded at https://atlas.fredhutch.org/nygc/multimodal-pbmc/. TCGA pan-cancer normalized RSEM gene expression data was downloaded from the GDC portal (https://portal.gdc.cancer.gov/). TCGA clinical information was retrieved from Liu et al.^78^.

## Code availability

The code for the experiments and visualizations can be found in the GitHub repository https://github.com/BoevaLab/ANS_supplementary_information. The Python package published in the GitHub repository https://github.com/lciernik/ANS_signature_scoring includes the implementations of all considered scoring methods. The repository also contains the R implementation of ANS.

## Acknowledgments

This work was supported by the Stiftung für angewandte Krebsforschung (SAKF) at the University Hospital Zurich (to A.K.), Swiss National Science Foundation (SNF) (projects CRSII5_209524 and 205321_207931 to F.B. and J.Y.), and the Hector Fellow Academy (L.C.).

## Author information

### Contributions

V.B., A.K., and L.C. conceived the study. J.Y. and F.B. preprocessed the cancer datasets and helped in the analysis and interpretation of results. L.C. developed the ANS scoring method and performed the analysis and interpretation of the benchmark and case study with the assistance of V.B. and A.K. A.K. conducted the association study of cancer EMT scores with histopathological types. L.C. wrote the paper with input from all authors.

### Corresponding author

Correspondence to <u>Valentina Boeva</u>. ETH Zurich, Universitatstrasse 6, 8092 Zurich, Switzerland

