## Supplementary Material for "ANS: Adjusted Neighborhood Scoring to improve gene signature-based cell annotation in single-cell RNA-seq data"

***Content***

**Supplementary Tables** 2

**Supplementary Figures** 11

**References** 22

### Supplementary Tables

**Table S1:** Gene signatures for each dataset used in the comparable score range experiment. https://docs.google.com/spreadsheets/d/1xsp9F4ob4w1se4Ln_0B05-b1E8ntbpnC/edit?usp=sharing&ouid=108568536620605288300&rtpof=true&sd=true

**Table S2.** Cell type annotation performance comparisons (balanced accuracy and f1-score weighted) based on ***non-overlapping*** signatures' scores and in the information of signatures scores computed with logistic regression.

| | | **Balanced Accuracy** | | | | | | | |
|---|---|---|---|---|---|---|---|---|---|
| | | **ANS** | **Jasmine_LH** | **Jasmine_OR** | **Scanpy** | **Seurat** | **Seurat_AG** | **Seurat_LVG** | **UCell** |
| **Hard labeling score** | BRCA (6 states) | **0.560** | 0.496 | 0.455 | 0.486 | 0.516 | 0.524 | 0.534 | 0.402 |
| | HGSOC (8 states) | **0.638** | 0.515 | 0.394 | 0.564 | 0.632 | 0.632 | 0.620 | 0.535 |
| | LUAD (3 states) | 0.897 | 0.885 | 0.791 | **0.914** | 0.894 | 0.896 | 0.825 | 0.839 |
| | cSCC (4 states) | **0.843** | 0.779 | 0.750 | 0.742 | 0.829 | 0.829 | 0.794 | 0.699 |
| | B, Mono-cytes, NK | 0.997 | 0.967 | 0.921 | **0.999** | 0.977 | 0.986 | 0.997 | **0.999** |
| | B-cell subtypes | **0.836** | 0.538 | 0.480 | 0.333 | 0.338 | 0.338 | 0.340 | 0.700 |
| | CD4 T-cell subtypes | **0.645** | 0.551 | 0.487 | 0.570 | 0.561 | 0.558 | 0.539 | 0.600 |
| | CD8 T-cell subtypes | **0.823** | 0.649 | 0.488 | 0.405 | 0.438 | 0.427 | 0.299 | 0.336 |
| **Information quantity (logistic regression 10-fold CV)** | BRCA (6 states) | 0.605 | 0.589 | 0.527 | 0.597 | 0.589 | 0.600 | 0.600 | **0.625** |
| | HGSOC (8 states) | 0.604 | 0.531 | 0.387 | **0.606** | 0.604 | 0.605 | 0.602 | 0.592 |
| | LUAD (3 states) | 0.892 | 0.870 | 0.811 | 0.878 | 0.882 | 0.890 | **0.923** | 0.899 |
| | cSCC (4 states) | 0.841 | 0.824 | 0.772 | **0.844** | 0.840 | 0.843 | 0.839 | 0.832 |
| | B, Mono-cytes, NK | 0.999 | 0.999 | 0.998 | 0.999 | 0.999 | 0.999 | **0.999** | 0.999 |
| | B-cell subtypes | **0.853** | 0.738 | 0.540 | 0.847 | **0.853** | 0.851 | 0.851 | 0.826 |
| | CD4 T-cell subtypes | 0.681 | 0.579 | 0.539 | 0.685 | 0.685 | **0.691** | 0.690 | 0.663 |
| | CD8 T-cell subtypes | **0.829** | 0.695 | 0.603 | 0.801 | 0.806 | 0.817 | 0.826 | 0.819 |

| | | F1 Score | | | | | | | |
|---|---|---|---|---|---|---|---|---|---|
| | | **ANS** | **Jasmine_LH** | **Jasmine_OR** | **Scanpy** | **Seurat** | **Seurat_AG** | **Seurat_LVG** | **UCell** |
| **Hard labeling score** | BRCA (6 states) | **0.492** | 0.453 | 0.407 | 0.426 | 0.457 | 0.464 | 0.472 | 0.309 |
| | HGSOC (8 states) | **0.599** | 0.547 | 0.492 | 0.513 | 0.591 | 0.587 | 0.555 | 0.552 |
| | LUAD (3 states) | 0.861 | 0.853 | 0.869 | 0.880 | 0.861 | 0.862 | 0.796 | **0.898** |
| | cSCC (4 states) | 0.825 | 0.800 | 0.751 | 0.784 | 0.830 | **0.832** | 0.820 | 0.759 |
| | B, Mono-cytes, NK | 0.996 | 0.984 | 0.962 | 0.999 | 0.958 | 0.975 | 0.996 | **0.999** |
| | B-cell subtypes | **0.834** | 0.474 | 0.345 | 0.104 | 0.115 | 0.117 | 0.125 | 0.757 |
| | CD4 T-cell subtypes | **0.457** | 0.216 | 0.255 | 0.354 | 0.362 | 0.361 | 0.352 | 0.448 |
| | CD8 T-cell subtypes | **0.825** | 0.504 | 0.248 | 0.398 | 0.484 | 0.460 | 0.308 | 0.306 |
| **Information quantity (logistic regression 10-fold CV)** | BRCA (6 states) | 0.598 | 0.588 | 0.528 | 0.586 | 0.581 | 0.592 | 0.591 | **0.618** |
| | HGSOC (8 states) | 0.717 | 0.661 | 0.575 | **0.718** | 0.716 | 0.717 | 0.714 | 0.709 |
| | LUAD (3 states) | 0.907 | 0.909 | 0.904 | 0.899 | 0.905 | 0.903 | **0.918** | 0.916 |
| | cSCC (4 states) | 0.869 | 0.857 | 0.824 | 0.869 | 0.868 | **0.870** | 0.867 | 0.861 |
| | B, Mono-cytes, NK | 0.999 | 0.999 | 0.999 | 0.999 | 0.999 | 0.999 | 0.999 | 0.999 |
| | B-cell subtypes | **0.894** | 0.799 | 0.618 | 0.890 | 0.893 | 0.892 | 0.893 | 0.874 |
| | CD4 T-cell subtypes | 0.731 | 0.603 | 0.562 | 0.741 | 0.745 | 0.746 | **0.747** | 0.716 |
| | CD8 T-cell subtypes | **0.913** | 0.819 | 0.734 | 0.906 | 0.909 | 0.909 | 0.907 | 0.896 |

**Table S3.** Cell type annotation performance comparisons (balanced accuracy and f1-score weighted) based on ***overlapping*** signatures' scores and in the information of signatures scores computed with logistic regression.

| | | Balanced Accuracy | | | | | | | |
|---|---|---|---|---|---|---|---|---|---|
| | | **ANS** | **Jasmine_LH** | **Jasmine_OR** | **Scanpy** | **Seurat** | **Seurat_AG** | **Seurat_LVG** | **UCell** |
| **Hard labeling score** | BRCA (6 states) | **0.575** | 0.522 | 0.515 | 0.519 | 0.533 | 0.536 | 0.543 | 0.411 |
| | HGSOC (8 states) | **0.646** | 0.570 | 0.418 | 0.577 | 0.637 | 0.637 | 0.626 | 0.553 |
| | LUAD (3 states) | 0.902 | 0.874 | 0.841 | **0.912** | 0.897 | 0.902 | 0.836 | 0.850 |
| | cSCC (4 states) | **0.846** | 0.785 | 0.757 | 0.742 | 0.822 | 0.827 | 0.792 | 0.696 |
| | B, Mono-cytes, NK | 0.998 | 0.997 | 0.992 | **0.999** | 0.988 | 0.993 | 0.998 | 0.998 |
| | B-cell subtypes | **0.850** | 0.667 | 0.442 | 0.333 | 0.412 | 0.378 | 0.413 | 0.679 |
| | CD4 T-cell subtypes | **0.669** | 0.585 | 0.581 | 0.167 | 0.189 | 0.198 | 0.170 | 0.208 |
| | CD8 T-cell subtypes | **0.784** | 0.732 | 0.722 | 0.447 | 0.390 | 0.376 | 0.438 | 0.415 |
| **Information quantity (logistic regression 10-fold CV)** | BRCA (6 states) | 0.612 | 0.608 | 0.576 | 0.602 | 0.605 | 0.611 | 0.606 | **0.636** |
| | HGSOC (8 states) | 0.612 | 0.537 | 0.395 | **0.613** | 0.612 | 0.612 | 0.607 | 0.600 |
| | LUAD (3 states) | 0.893 | 0.856 | 0.860 | 0.881 | 0.882 | 0.882 | **0.914** | 0.913 |
| | cSCC (4 states) | 0.840 | 0.825 | 0.779 | **0.842** | 0.839 | 0.842 | 0.839 | 0.834 |
| | B, Mono-cytes, NK | 0.999 | 0.999 | 0.998 | **0.999** | 0.999 | 0.999 | 0.999 | 0.999 |
| | B-cell subtypes | 0.884 | 0.726 | 0.457 | 0.865 | **0.885** | 0.883 | 0.881 | 0.857 |
| | CD4 T-cell subtypes | 0.760 | 0.624 | 0.570 | 0.728 | 0.757 | **0.762** | 0.750 | 0.734 |
| | CD8 T-cell subtypes | 0.864 | 0.762 | 0.719 | 0.847 | 0.866 | 0.869 | **0.875** | 0.854 |

| | | F1 Score | | | | | | | |
|---|---|---|---|---|---|---|---|---|---|
| | | **ANS** | **Jasmine_LH** | **Jasmine_OR** | **Scanpy** | **Seurat** | **Seurat_AG** | **Seurat_LVG** | **UCell** |
| **Hard labeling score** | BRCA (6 states) | **0.503** | 0.470 | 0.458 | 0.447 | 0.473 | 0.475 | 0.479 | 0.326 |
| | HGSOC (8 states) | **0.600** | 0.555 | 0.485 | 0.477 | 0.584 | 0.585 | 0.537 | 0.511 |
| | LUAD (3 states) | 0.866 | 0.847 | 0.872 | 0.881 | 0.864 | 0.868 | 0.809 | **0.902** |
| | cSCC (4 states) | 0.827 | 0.806 | 0.763 | 0.783 | 0.830 | **0.832** | 0.819 | 0.753 |
| | B, Mono-cytes, NK | 0.997 | 0.998 | 0.996 | **0.999** | 0.977 | 0.987 | 0.998 | 0.999 |
| | B-cell subtypes | **0.869** | 0.667 | 0.362 | 0.104 | 0.331 | 0.243 | 0.334 | 0.758 |
| | CD4 T-cell subtypes | **0.451** | 0.319 | 0.340 | 0.008 | 0.020 | 0.053 | 0.009 | 0.157 |
| | CD8 T-cell subtypes | **0.851** | 0.753 | 0.756 | 0.455 | 0.389 | 0.371 | 0.453 | 0.417 |
| **Information quantity (logistic regression 10-fold CV)** | BRCA (6 states) | 0.606 | 0.605 | 0.571 | 0.594 | 0.599 | 0.605 | 0.600 | **0.629** |
| | HGSOC (8 states) | 0.726 | 0.673 | 0.585 | **0.727** | 0.726 | 0.726 | 0.721 | 0.719 |
| | LUAD (3 states) | 0.909 | 0.912 | 0.908 | 0.903 | 0.904 | 0.904 | 0.917 | **0.918** |
| | cSCC (4 states) | 0.867 | 0.858 | 0.830 | 0.868 | 0.868 | **0.870** | 0.866 | 0.864 |
| | B, Mono-cytes, NK | 1.000 | 1.000 | 0.999 | **1.000** | 1.000 | 1.000 | 0.999 | 0.999 |
| | B-cell subtypes | 0.916 | 0.794 | 0.563 | 0.903 | **0.918** | 0.916 | 0.915 | 0.899 |
| | CD4 T-cell subtypes | 0.808 | 0.672 | 0.610 | 0.782 | 0.806 | **0.810** | 0.803 | 0.793 |
| | CD8 T-cell subtypes | **0.935** | 0.871 | 0.846 | 0.934 | 0.931 | 0.934 | 0.934 | 0.930 |

**Table S4.** Full ESCC and LUAD cancer EMT signature genes and per gene information: ESCC- and LUAD-specific cancer EMT signature (https://docs.google.com/spreadsheets/d/1-NRBqLv2TIfipSGged5yICR84n3XXc1x/edit?usp=sharing&ouid=108568536620605288300&rtpof=true&sd=true ). For each gene, the table indicates if it has been included in any considered pan-cancer EMT signature, in EMTome[1], in dbEMT[2], or if we have found literature associating the gene with EMT.

**Table S5.** AUCPRC values measuring the performance of signature scores to discriminate malignant EMT cells, CAFs, malignant cells not undergoing EMT, and all other cell types using the LUAD- and ESCC-specific cancer EMT and the pan-cancer EMT signatures for scoring.

| | **AUCPRC, cancer MES-like *vs.* CAFs** | | | | **AUCPRC, cancer MES-like *vs.* other cancer cells** | | | | **AUCPRC, cancer MES-like *vs.* rest** | | | |
|---|---|---|---|---|---|---|---|---|---|---|---|---|
| | **ESCC** | **LUAD** | **CRC** | **BRCA** | **ESCC** | **LUAD** | **CRC** | **BRCA** | **ESCC** | **LUAD** | **CRC** | **BRCA** |
| **ESCC and LUAD cancer EMT signature (this paper)** | **0,952** | **0,988** | **0,987** | **0,907** | 0,595 | 0,545 | 0,705 | 0,702 | **0,592** | **0,513** | 0,701 | **0,511** |
| **Hallmark EMT signature**[3] | 0,065 | 0,333 | 0,442 | 0,460 | **0,684** | **0,596** | 0,631 | **0,860** | 0,038 | 0,049 | 0,042 | 0,172 |
| **pEMT gene module**[4] | 0,428 | 0,523 | 0,651 | 0,743 | 0,520 | 0,474 | 0,563 | 0,708 | 0,332 | 0,185 | 0,277 | 0,437 |
| **EMT signature Foroutan *et al.***[5] | 0,065 | 0,382 | 0,445 | 0,459 | 0,368 | 0,451 | **0,850** | 0,825 | 0,033 | 0,128 | 0,190 | 0,194 |
| **EMT signature Groeger *et al.***[6] | 0,065 | 0,333 | 0,442 | 0,456 | 0,607 | 0,413 | 0,224 | 0,744 | 0,024 | 0,014 | 0,019 | 0,115 |
| **EMT signature Hollern *et al.***[7] | 0,070 | 0,589 | 0,880 | 0,472 | 0,104 | 0,581 | 0,795 | 0,600 | 0,028 | 0,245 | **0,707** | 0,106 |
| **EMT signature *Mak et al.***[8] | 0,065 | 0,334 | 0,442 | 0,463 | 0,418 | 0,514 | 0,566 | 0,794 | 0,026 | 0,067 | 0,085 | 0,182 |
| **EMT signature Tan *et al.***[9] | 0,065 | 0,411 | 0,813 | 0,454 | 0,137 | 0,520 | 0,807 | 0,610 | 0,031 | 0,218 | 0,688 | 0,196 |

**Table S6**. Association of ESCC- and LUAD-specific cancer EMT signature scores and histological subtypes in TCGA (only cancer types with at least 1 histotype are included).

| Cancer type | P-value | P-value adjusted |
|---|---|---|
| ACC | 0.428860685 | 0.514633 |
| BRCA | 0.002046152 | 0.00568 |
| CESC | 0.000635924 | 0.00259 |
| CHOL | 0.953345618 | 0.953346 |
| COAD | 0.476786946 | 0.520131 |
| DLBC | 0.370744213 | 0.49751 |
| ESCA | 0.001724259 | 0.00568 |
| GBM | 0.47320919 | 0.520131 |
| HNSC | 0.00019615 | 0.001177 |
| LGG | 0.011776278 | 0.028263 |
| LIHC | 0.000647502 | 0.00259 |
| MESO | 0.126606329 | 0.233735 |
| PAAD | 0.002129934 | 0.00568 |
| PCPG | 0.37313238 | 0.49751 |
| PRAD | 0.311602561 | 0.467404 |
| READ | 0.784831706 | 0.818955 |
| SARC | 0.070232513 | 0.140465 |
| STAD | 0.039594271 | 0.086388 |
| TGCT | 3.15932E-07 | 2.53E-06 |
| THCA | 8.74626E-33 | 2.1E-31 |
| THYM | 5.50201E-09 | 6.6E-08 |
| UCEC | 0.420181429 | 0.514633 |
| UCS | 0.136490018 | 0.233983 |
| UVM | 0.296780396 | 0.467404 |

### Supplementary Figures

**Figure S1:** Sorted average gene expression values of CRC with expression bins for Tirosh et al.[10] based gene signature scoring methods.

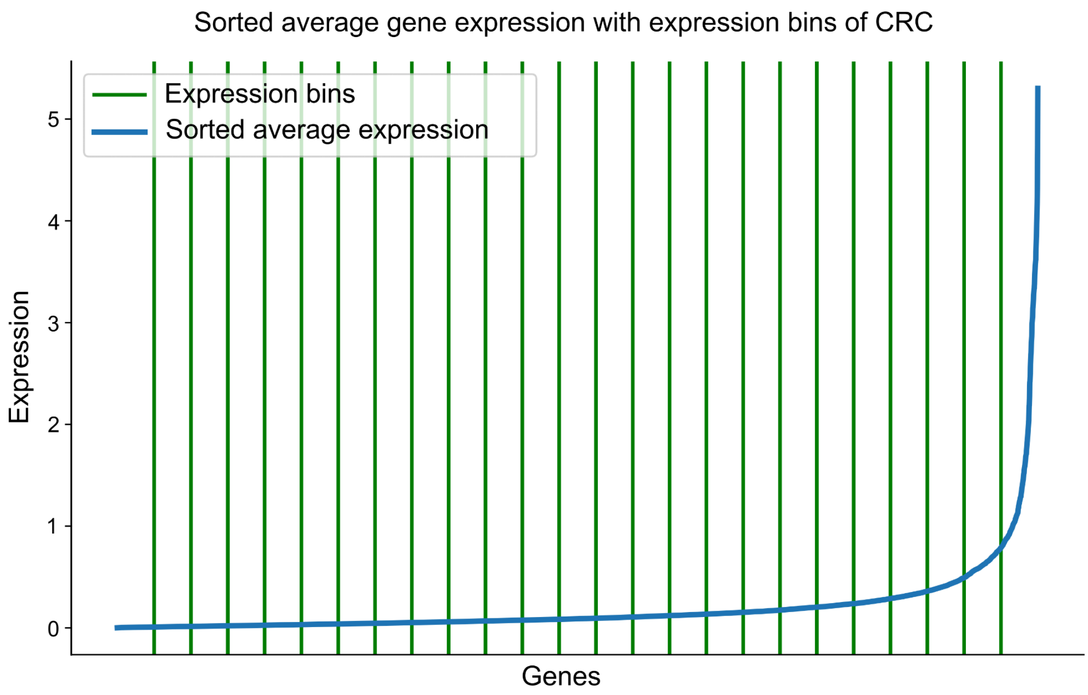

**Figure S2**: Benchmarking computation times of scoring methods. This figure presents the computation times required for scoring across varying dataset sizes, with four distinct signature lengths (1, 10, 100, 1000).

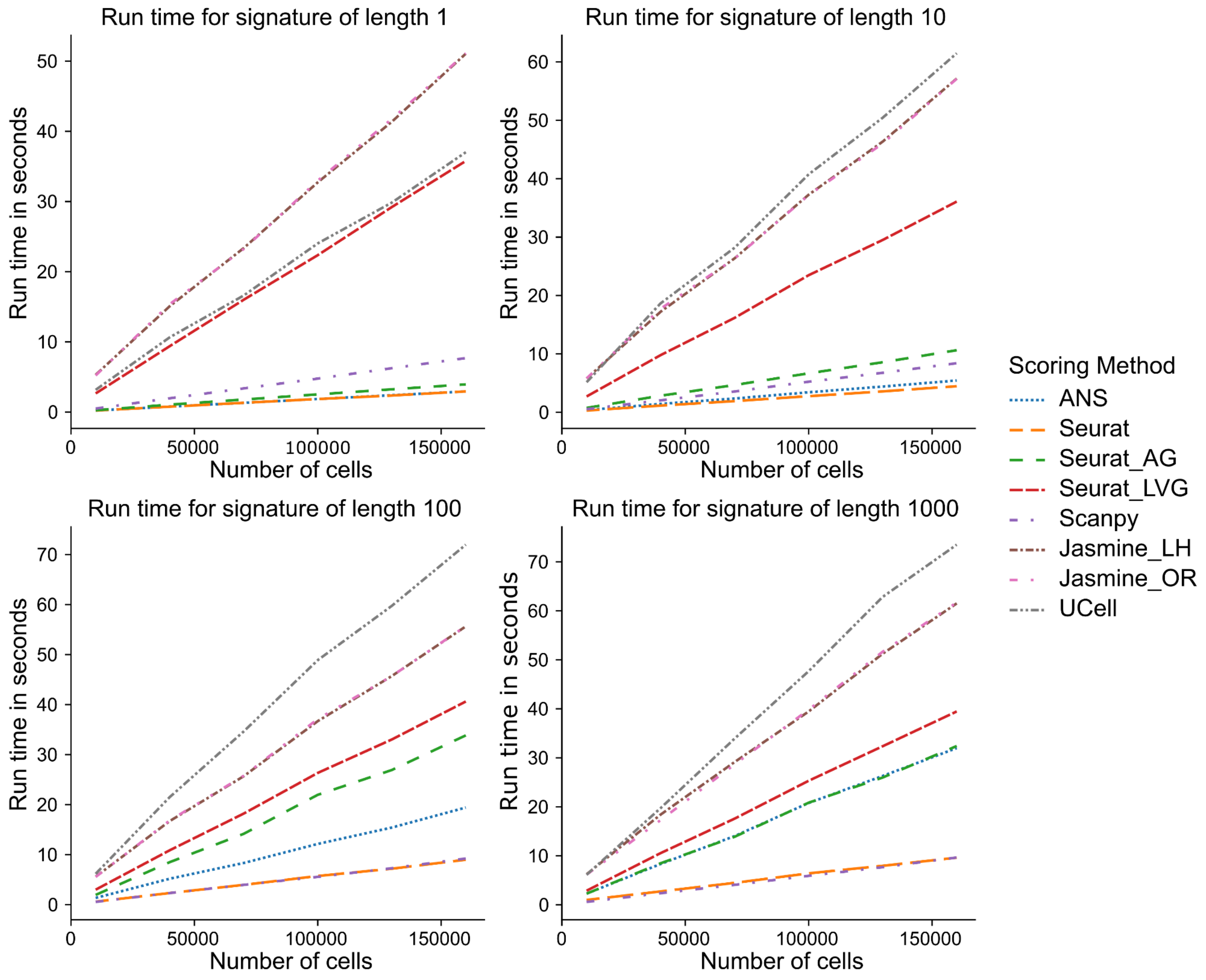

**Figure S3:** Control genes selection bias in Tirosh-based methods for the top 8% of highly expressed genes in B memory kappa, CD8 TEM 2, CD14 Monocytes, and NK 3 cells within the PBMC dataset. The x-axis illustrates genes sorted by their average expression levels, while the y-axis depicts the mean and standard deviation of the scores across all cells for a single-gene signature. Vertical dashed lines indicate the expression bin boundary and the top 50 highly expressed genes. The bias of a scoring method is indicated by how far the mean score of a gene deviates from zero.

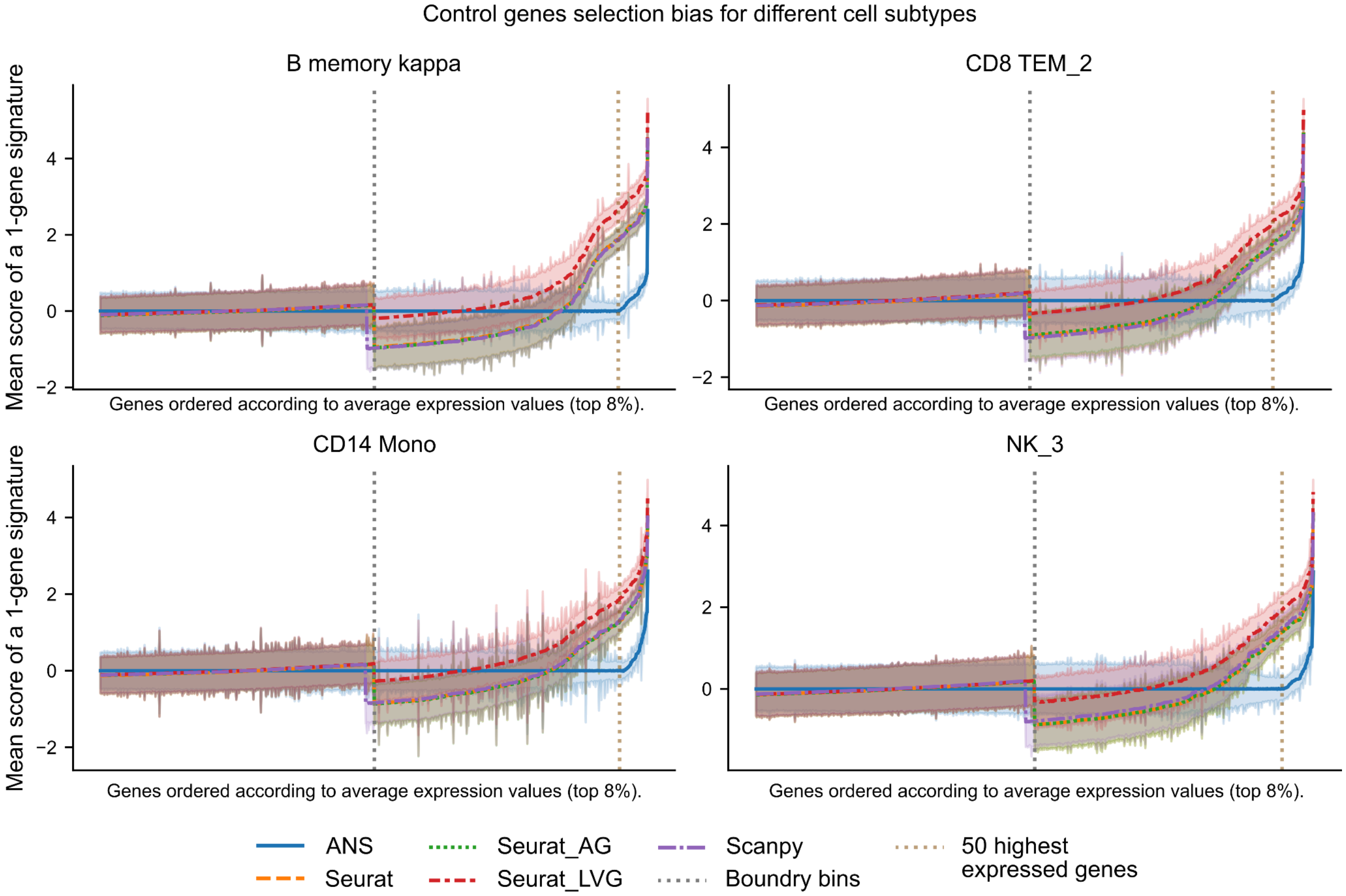

**Figure S4**: Benchmark on ESCC **a.** The influence of dataset composition on scoring ESCC cells using a 100-gene signature associated with malignant cells. Each dot represents the mean score for all cells within a sample, grouped by malignancy (malignant in blue or non-malignant in orange) and scoring mode (scoring all the samples together or individually). A black horizontal bar represents the mean value of all dots within each group. We observed a decrease in score variance when scoring on all samples. **b.** The minimum required signature length per scoring method for perfect classification (AUCROC of 1) when discriminating between malignant and non-malignant cell scores in ESCC. **c.** The robustness to noise in a signature when discriminating between malignant and non-malignant cells in ESCC. Starting with a 100-gene signature, we performed iterative replacements of genes with random genes that exhibited a |log2FC| < 0.5 and an adjusted p-value > 0.1 during differential gene expression (DGEX) analysis between malignant and non-malignant cells. The standard variation was calculated from 20 simulation runs.

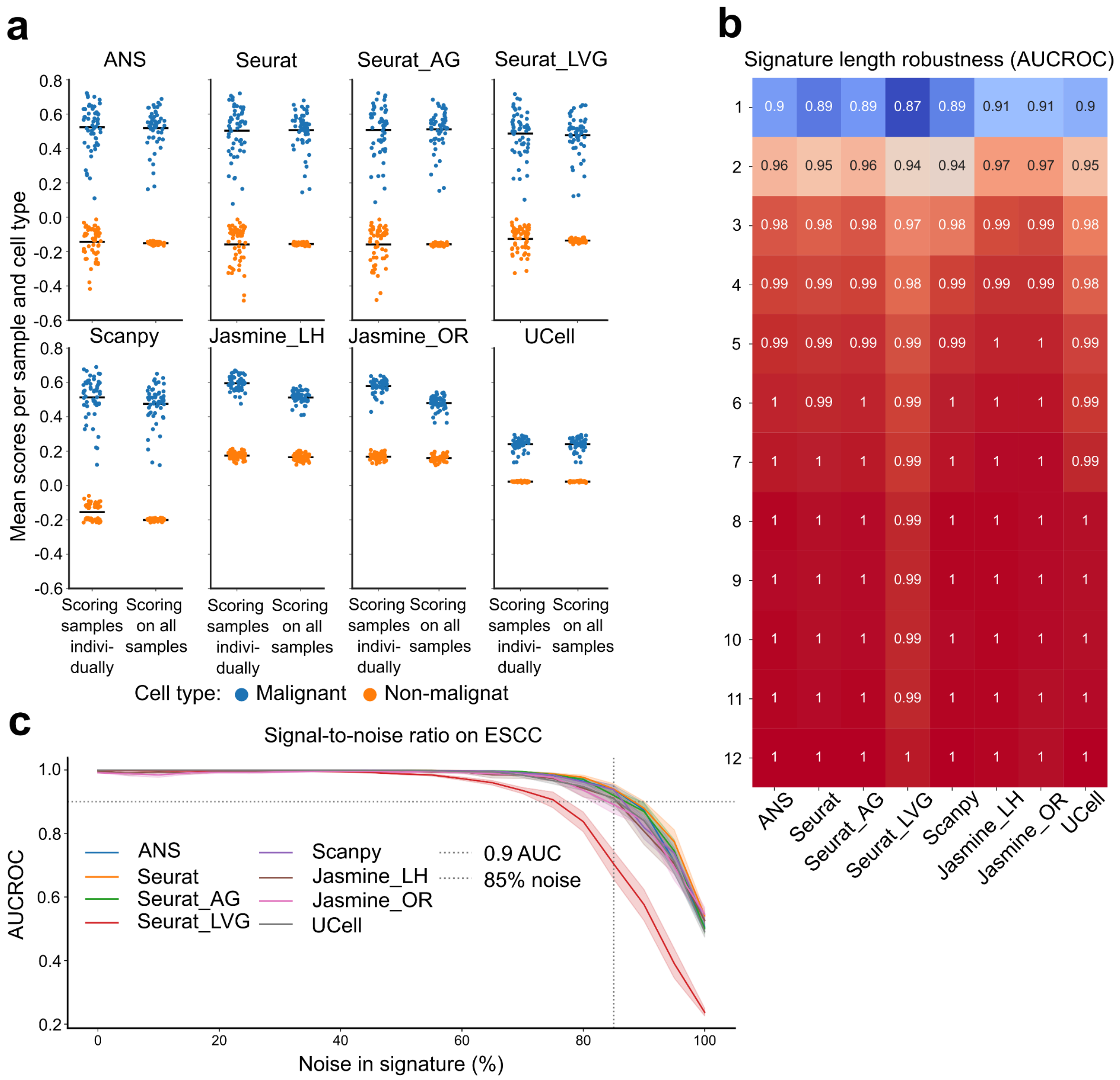

**Figure S5:** Score variances comparison between scoring modes (scoring sample individually versus scoring the entire dataset) for all scoring methods and two malignancy types on CRC and ESCC.

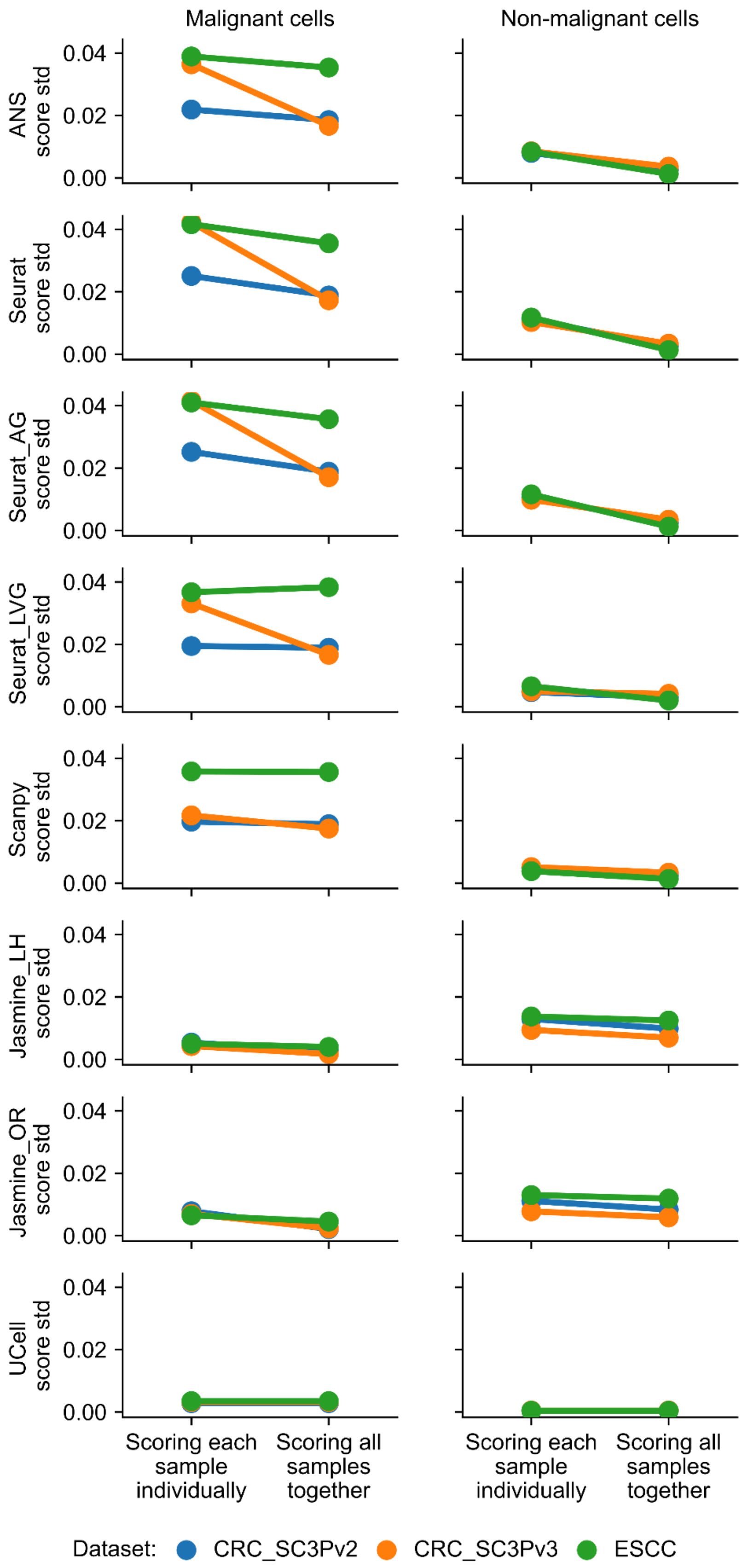

**Figure S6:** Sensitivity to signature length and signal-to-noise ratio in CRC. **a.** The minimum required signature length per scoring method for perfect classification (AUCROC of 1) when discriminating between malignant and non-malignant cell scores in CRC. **b.** The robustness to noise in a signature when discriminating between malignant and non-malignant cells in CRC. Starting with a 100-gene signature, we performed iterative replacements of genes with random genes that exhibited a |log2FC| < 0.5 and an adjusted p-value > 0.1 during differential gene expression (DGEX) analysis between malignant and non-malignant cells. The standard variation was calculated from 20 simulation runs.

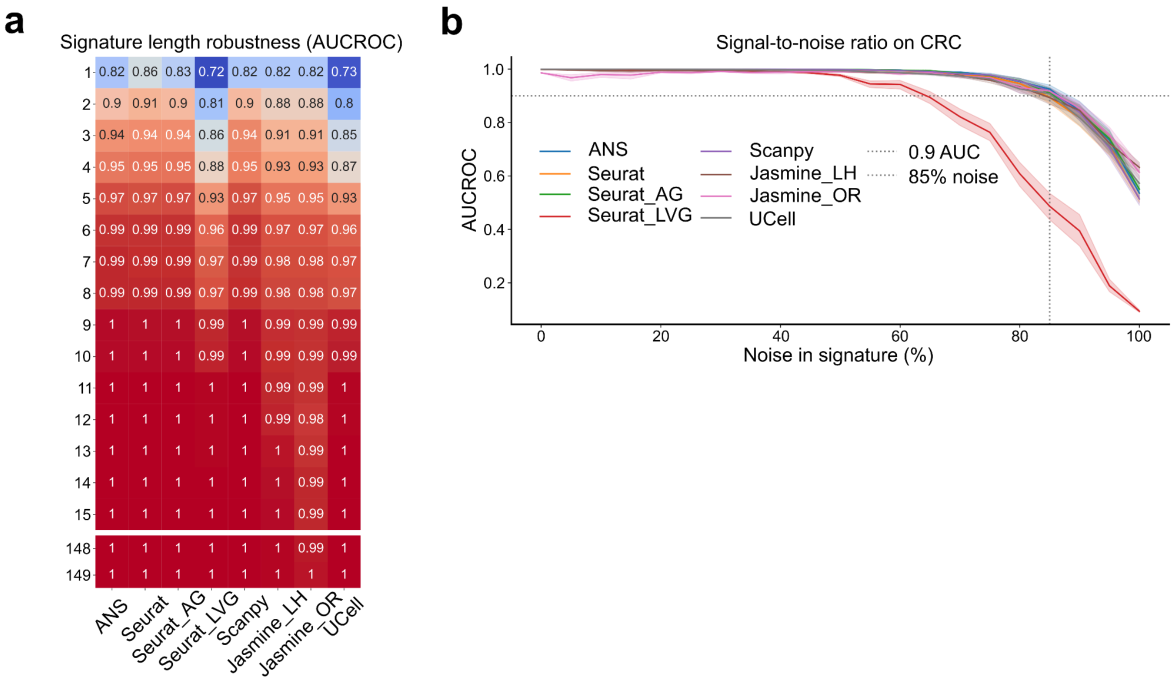

**Figure S7:** Confusion matrices for cell type/state prediction across multiple methods and datasets. **a**, Comparison of prediction performance between ANS, Jasmine_LH, Jasmine_OR, and Scanpy for cancer datasets (BRCA, LUAD, HGSOC, cSCC). **b**, Results for Seurat, Seurat_AG, Seurat_LVG, and UCell on the same cancer datasets. **c-d**, Performance evaluation across immune cell subsets using the same methods, showing B/Monocytes/NK cells, B-cell subtypes, CD4 T-cell subtypes, and CD8 T-cell subtypes. Each matrix shows the relationship between true (y-axis) and predicted (x-axis) cell states, with values summing to one per row. Color scale ranges from blue (0) to red (1), with balanced accuracy (bal. acc.) shown for each matrix. **Non-overlapping** gene signatures were used to score cell states/types in each dataset.

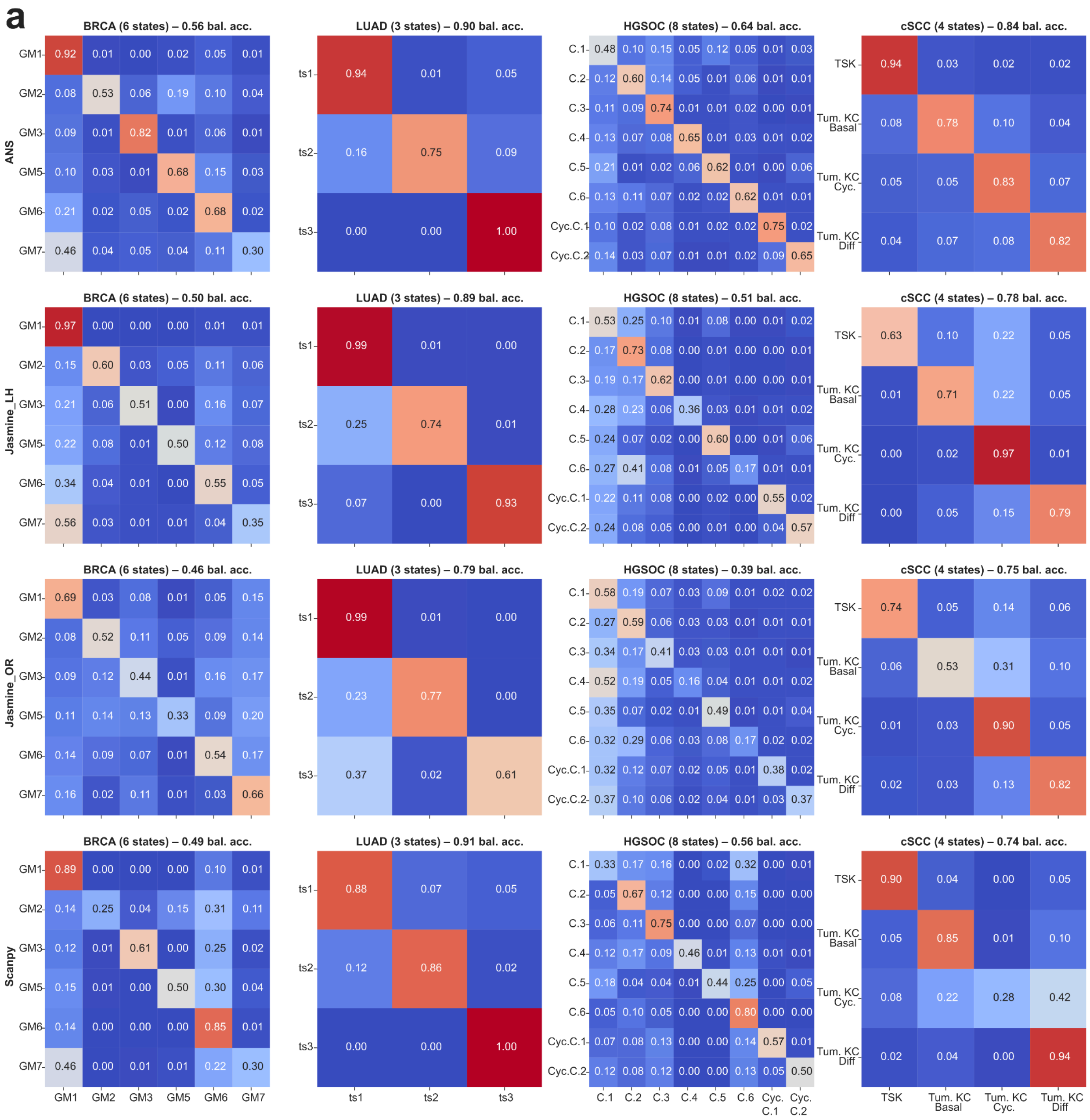

**b**

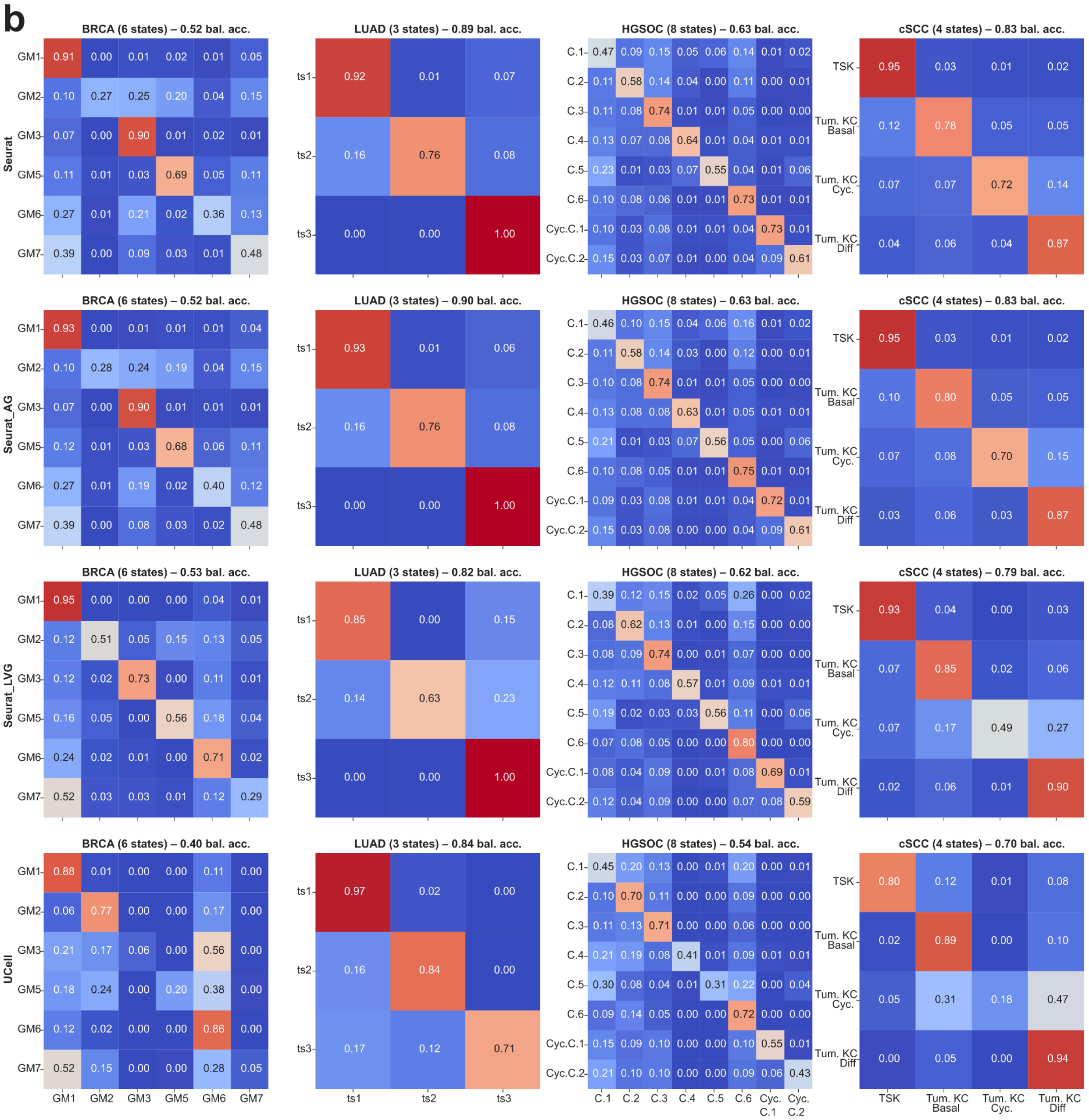
BRCA (6 states) – 0.52 bal. acc.
LUAD (3 states) – 0.89 bal. acc.
HGSOC (8 states) – 0.63 bal. acc.
cSCC (4 states) – 0.83 bal. acc.
BRCA (6 states) – 0.52 bal. acc.
LUAD (3 states) – 0.90 bal. acc.
HGSOC (8 states) – 0.63 bal. acc.
cSCC (4 states) – 0.83 bal. acc.
BRCA (6 states) – 0.53 bal. acc.
LUAD (3 states) – 0.82 bal. acc.
HGSOC (8 states) – 0.62 bal. acc.
cSCC (4 states) – 0.79 bal. acc.
BRCA (6 states) – 0.40 bal. acc.
LUAD (3 states) – 0.84 bal. acc.
HGSOC (8 states) – 0.54 bal. acc.
cSCC (4 states) – 0.70 bal. acc.
Seurat
Seurat_AG
Seurat_LVG
UCell
GM1 GM2 GM3 GM5 GM6 GM7
ts1 ts2 ts3
C.1 C.2 C.3 C.4 C.5 C.6 Cyc.C.1 Cyc.C.2
TSK
Tum. KC Basal
Tum. KC Cyc.
Tum. KC Diff

c

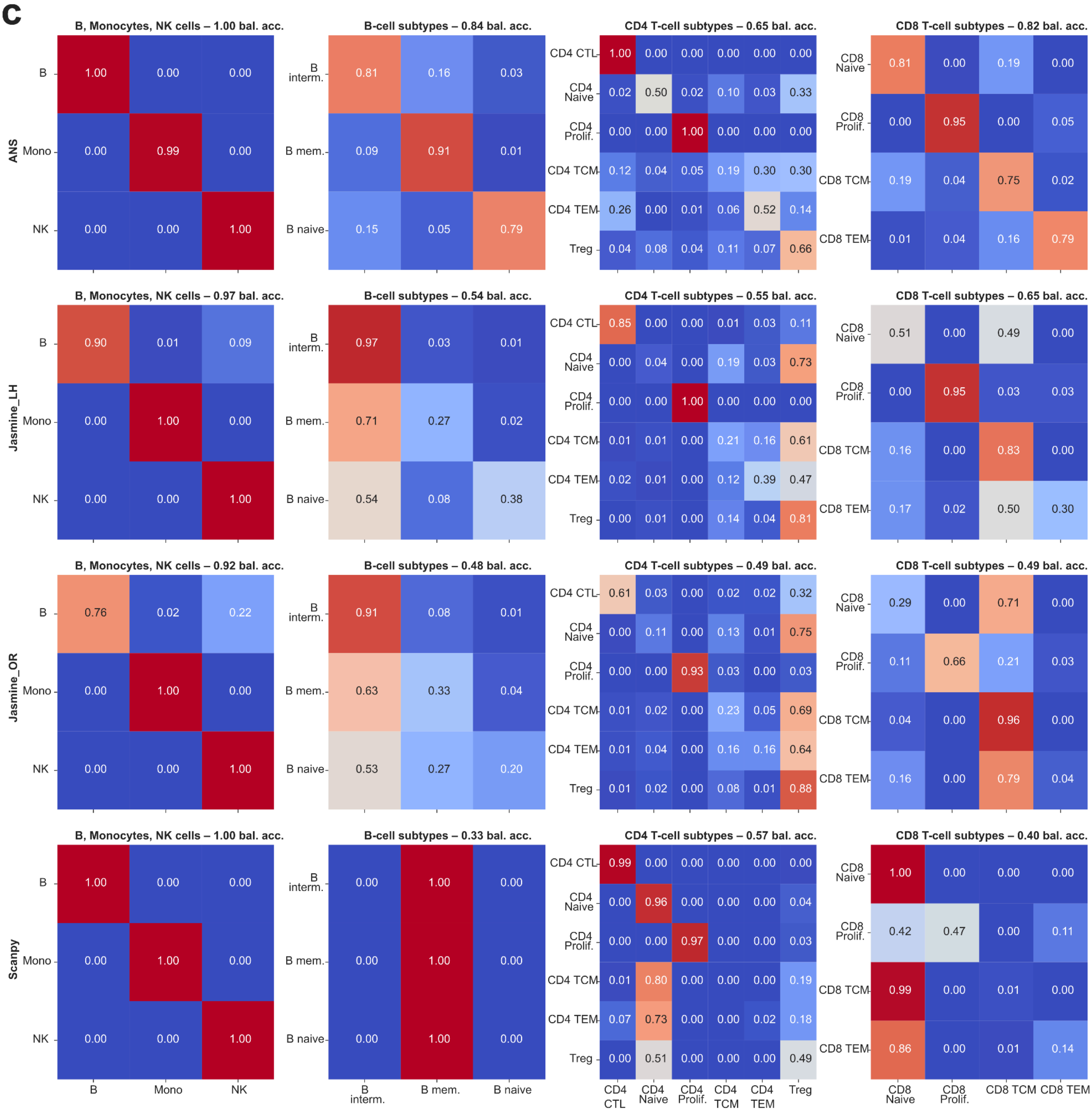
B, Monocytes, NK cells – 1.00 bal. acc.
B-cell subtypes – 0.84 bal. acc.
CD4 T-cell subtypes – 0.65 bal. acc.
CD8 T-cell subtypes – 0.82 bal. acc.
B, Monocytes, NK cells – 0.97 bal. acc.
B-cell subtypes – 0.54 bal. acc.
CD4 T-cell subtypes – 0.55 bal. acc.
CD8 T-cell subtypes – 0.65 bal. acc.
B, Monocytes, NK cells – 0.92 bal. acc.
B-cell subtypes – 0.48 bal. acc.
CD4 T-cell subtypes – 0.49 bal. acc.
CD8 T-cell subtypes – 0.49 bal. acc.
B, Monocytes, NK cells – 1.00 bal. acc.
B-cell subtypes – 0.33 bal. acc.
CD4 T-cell subtypes – 0.57 bal. acc.
CD8 T-cell subtypes – 0.40 bal. acc.
ANS
Jasmine_LH
Jasmine_OR
Scanpy
B
Mono
NK
B interm.
B mem.
B naive
CD4 CTL
CD4 Naive
CD4 Prolif.
CD4 TCM
CD4 TEM
Treg
CD8 Naive
CD8 Prolif.
CD8 TCM
CD8 TEM

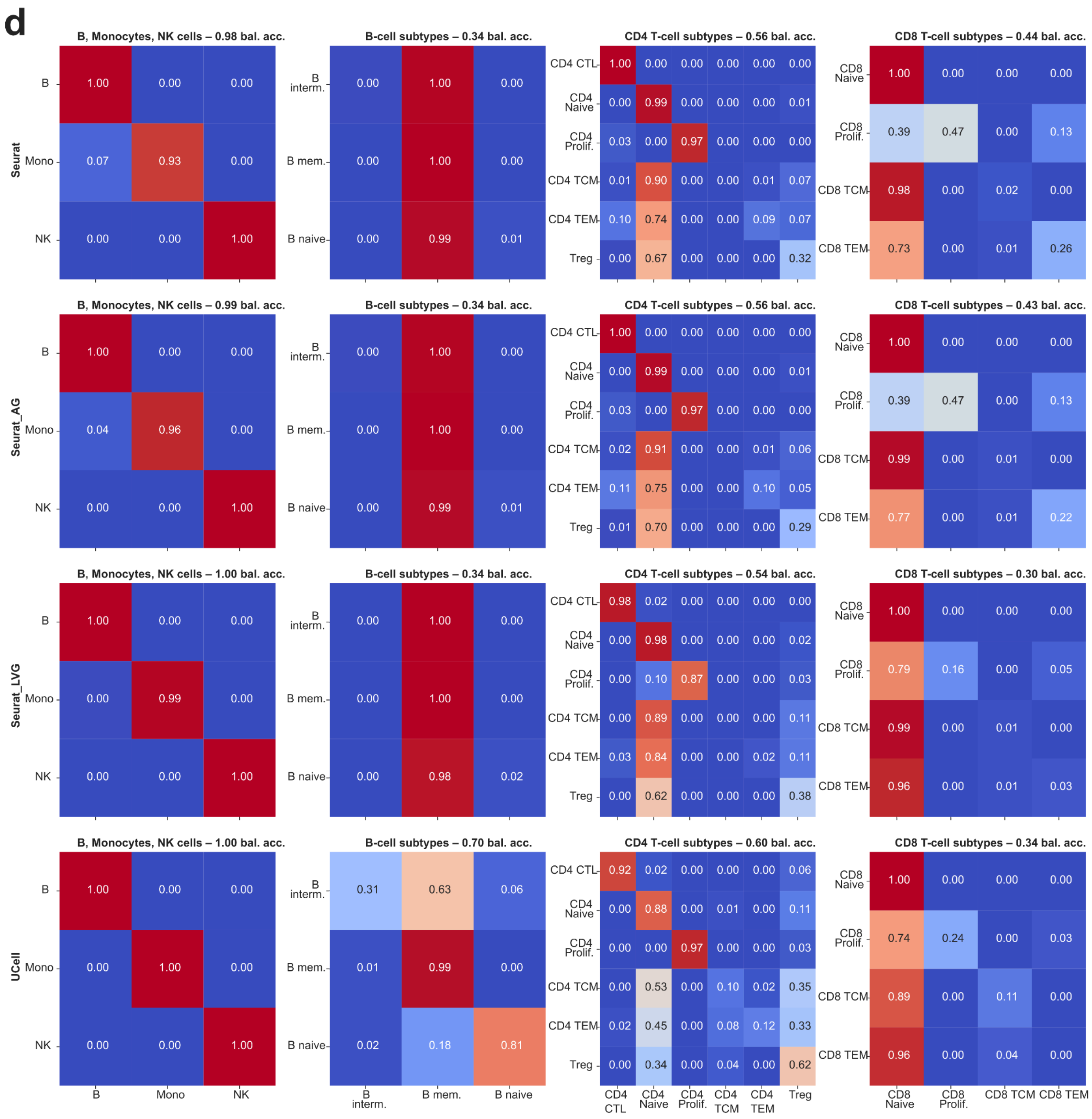
d
Seurat
Seurat_AG
Seurat_LVG
UCell
B, Monocytes, NK cells – 0.98 bal. acc.
B 1.00 0.00 0.00
Mono 0.07 0.93 0.00
NK 0.00 0.00 1.00
B-cell subtypes – 0.34 bal. acc.
B interm. 0.00 1.00 0.00
B mem. 0.00 1.00 0.00
B naive 0.00 0.99 0.01
CD4 T-cell subtypes – 0.56 bal. acc.
CD4 CTL 1.00 0.00 0.00 0.00 0.00 0.00
CD4 Naive 0.00 0.99 0.00 0.00 0.00 0.01
CD4 Prolif. 0.03 0.00 0.97 0.00 0.00 0.00
CD4 TCM 0.01 0.90 0.00 0.00 0.01 0.07
CD4 TEM 0.10 0.74 0.00 0.00 0.09 0.07
Treg 0.00 0.67 0.00 0.00 0.00 0.32
CD8 T-cell subtypes – 0.44 bal. acc.
CD8 Naive 1.00 0.00 0.00 0.00
CD8 Prolif. 0.39 0.47 0.00 0.13
CD8 TCM 0.98 0.00 0.02 0.00
CD8 TEM 0.73 0.00 0.01 0.26
B, Monocytes, NK cells – 0.99 bal. acc.
B 1.00 0.00 0.00
Mono 0.04 0.96 0.00
NK 0.00 0.00 1.00
B-cell subtypes – 0.34 bal. acc.
B interm. 0.00 1.00 0.00
B mem. 0.00 1.00 0.00
B naive 0.00 0.99 0.01
CD4 T-cell subtypes – 0.56 bal. acc.
CD4 CTL 1.00 0.00 0.00 0.00 0.00 0.00
CD4 Naive 0.00 0.99 0.00 0.00 0.00 0.01
CD4 Prolif. 0.03 0.00 0.97 0.00 0.00 0.00
CD4 TCM 0.02 0.91 0.00 0.00 0.01 0.06
CD4 TEM 0.11 0.75 0.00 0.00 0.10 0.05
Treg 0.01 0.70 0.00 0.00 0.00 0.29
CD8 T-cell subtypes – 0.43 bal. acc.
CD8 Naive 1.00 0.00 0.00 0.00
CD8 Prolif. 0.39 0.47 0.00 0.13
CD8 TCM 0.99 0.00 0.01 0.00
CD8 TEM 0.77 0.00 0.01 0.22
B, Monocytes, NK cells – 1.00 bal. acc.
B 1.00 0.00 0.00
Mono 0.00 0.99 0.00
NK 0.00 0.00 1.00
B-cell subtypes – 0.34 bal. acc.
B interm. 0.00 1.00 0.00
B mem. 0.00 1.00 0.00
B naive 0.00 0.98 0.02
CD4 T-cell subtypes – 0.54 bal. acc.
CD4 CTL 0.98 0.02 0.00 0.00 0.00 0.00
CD4 Naive 0.00 0.98 0.00 0.00 0.00 0.02
CD4 Prolif. 0.00 0.10 0.87 0.00 0.00 0.03
CD4 TCM 0.00 0.89 0.00 0.00 0.00 0.11
CD4 TEM 0.03 0.84 0.00 0.00 0.02 0.11
Treg 0.00 0.62 0.00 0.00 0.00 0.38
CD8 T-cell subtypes – 0.30 bal. acc.
CD8 Naive 1.00 0.00 0.00 0.00
CD8 Prolif. 0.79 0.16 0.00 0.05
CD8 TCM 0.99 0.00 0.01 0.00
CD8 TEM 0.96 0.00 0.01 0.03
B, Monocytes, NK cells – 1.00 bal. acc.
B 1.00 0.00 0.00
Mono 0.00 1.00 0.00
NK 0.00 0.00 1.00
B-cell subtypes – 0.70 bal. acc.
B interm. 0.31 0.63 0.06
B mem. 0.01 0.99 0.00
B naive 0.02 0.18 0.81
CD4 T-cell subtypes – 0.60 bal. acc.
CD4 CTL 0.92 0.02 0.00 0.00 0.00 0.06
CD4 Naive 0.00 0.88 0.00 0.01 0.00 0.11
CD4 Prolif. 0.00 0.00 0.97 0.00 0.00 0.03
CD4 TCM 0.00 0.53 0.00 0.10 0.02 0.35
CD4 TEM 0.02 0.45 0.00 0.08 0.12 0.33
Treg 0.00 0.34 0.00 0.04 0.00 0.62
CD8 T-cell subtypes – 0.34 bal. acc.
CD8 Naive 1.00 0.00 0.00 0.00
CD8 Prolif. 0.74 0.24 0.00 0.03
CD8 TCM 0.89 0.00 0.11 0.00
CD8 TEM 0.96 0.00 0.04 0.00
B Mono NK
B interm. B mem. B naive
CD4 CTL CD4 Naive CD4 Prolif. CD4 TCM CD4 TEM Treg
CD8 Naive CD8 Prolif. CD8 TCM CD8 TEM

**Figure S8:** Violin plots showing the distribution of scores for each scored signature per dataset. For each cell state or type, we expect the highest scores for the signature associated with this state or type (matching dark and light shades). **a**, Breast cancer (BRCA). **b**, Lung adenocarcinoma (LUAD). **c**, High-grade serous ovarian cancer (HGSOC). **d**, Squamous cell carcinoma (SCC). **e**, CD4+ T cell subtypes. **f**, CD8+ T cell subtypes.

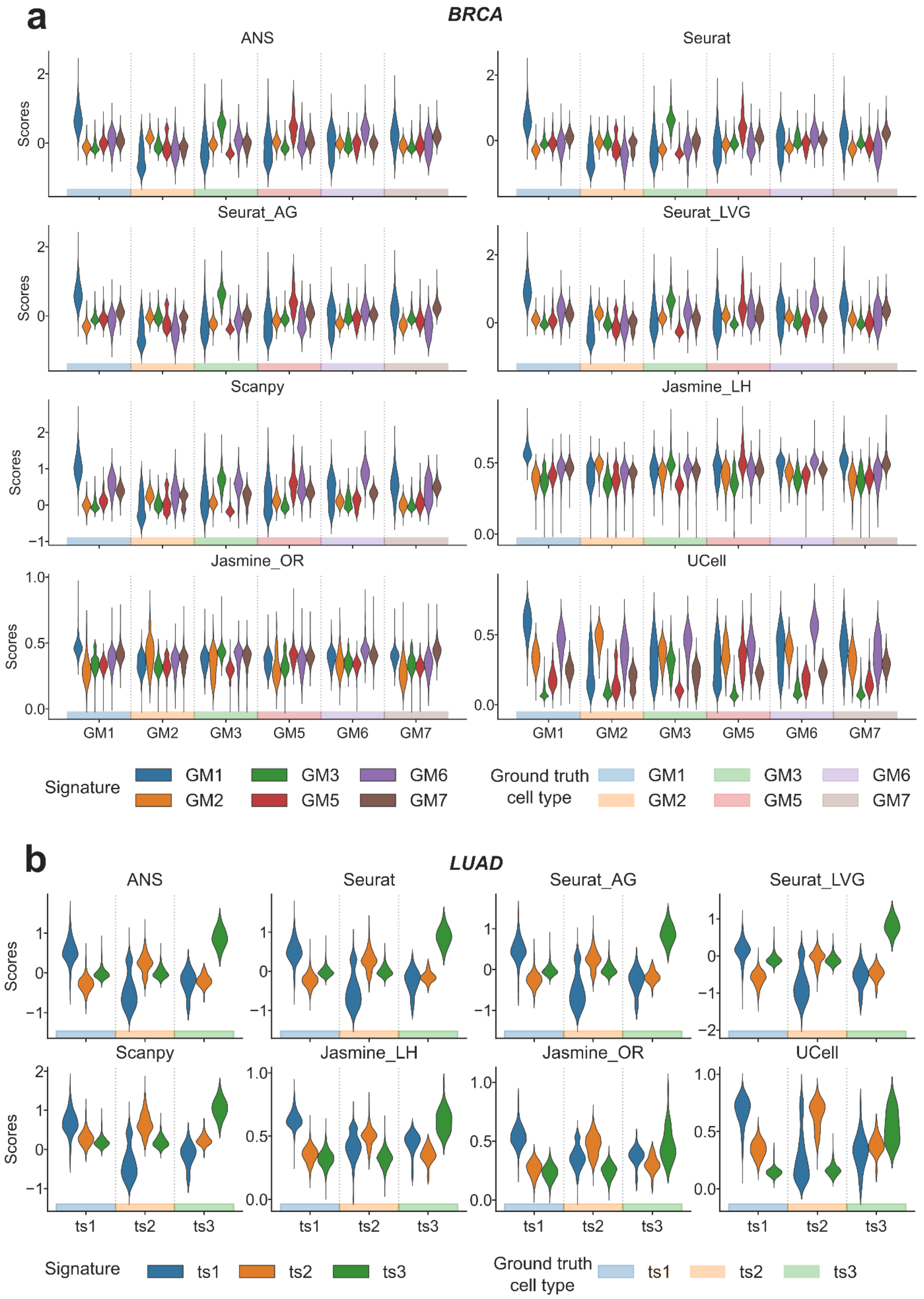

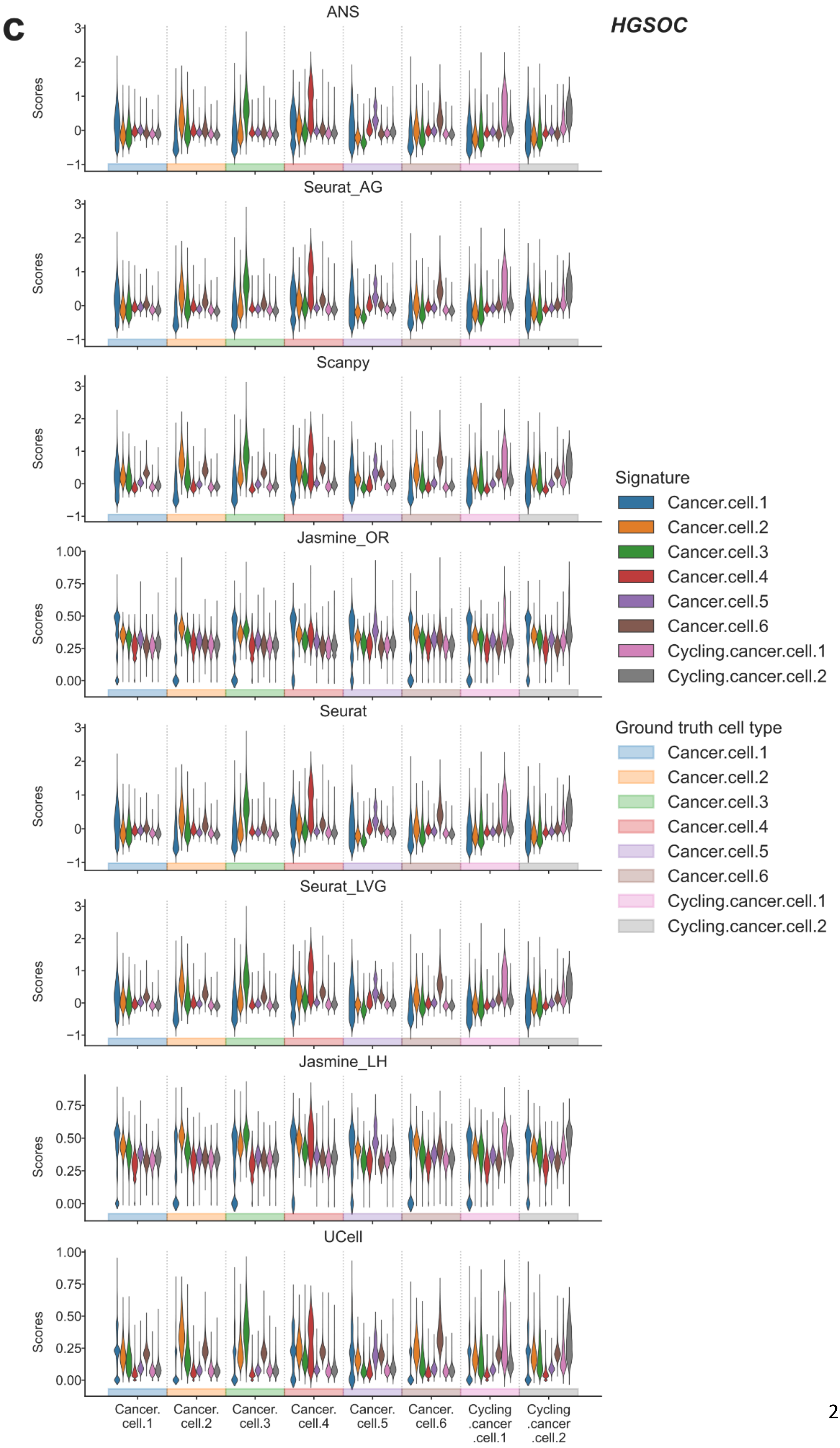
c
HGSOC
ANS
Seurat_AG
Scanpy
Jasmine_OR
Seurat
Seurat_LVG
Jasmine_LH
UCell
Scores
Signature
Cancer.cell.1
Cancer.cell.2
Cancer.cell.3
Cancer.cell.4
Cancer.cell.5
Cancer.cell.6
Cycling.cancer.cell.1
Cycling.cancer.cell.2
Ground truth cell type
Cancer.cell.1
Cancer.cell.2
Cancer.cell.3
Cancer.cell.4
Cancer.cell.5
Cancer.cell.6
Cycling.cancer.cell.1
Cycling.cancer.cell.2
Cancer. cell.1
Cancer. cell.2
Cancer. cell.3
Cancer. cell.4
Cancer. cell.5
Cancer. cell.6
Cycling .cancer .cell.1
Cycling .cancer .cell.2

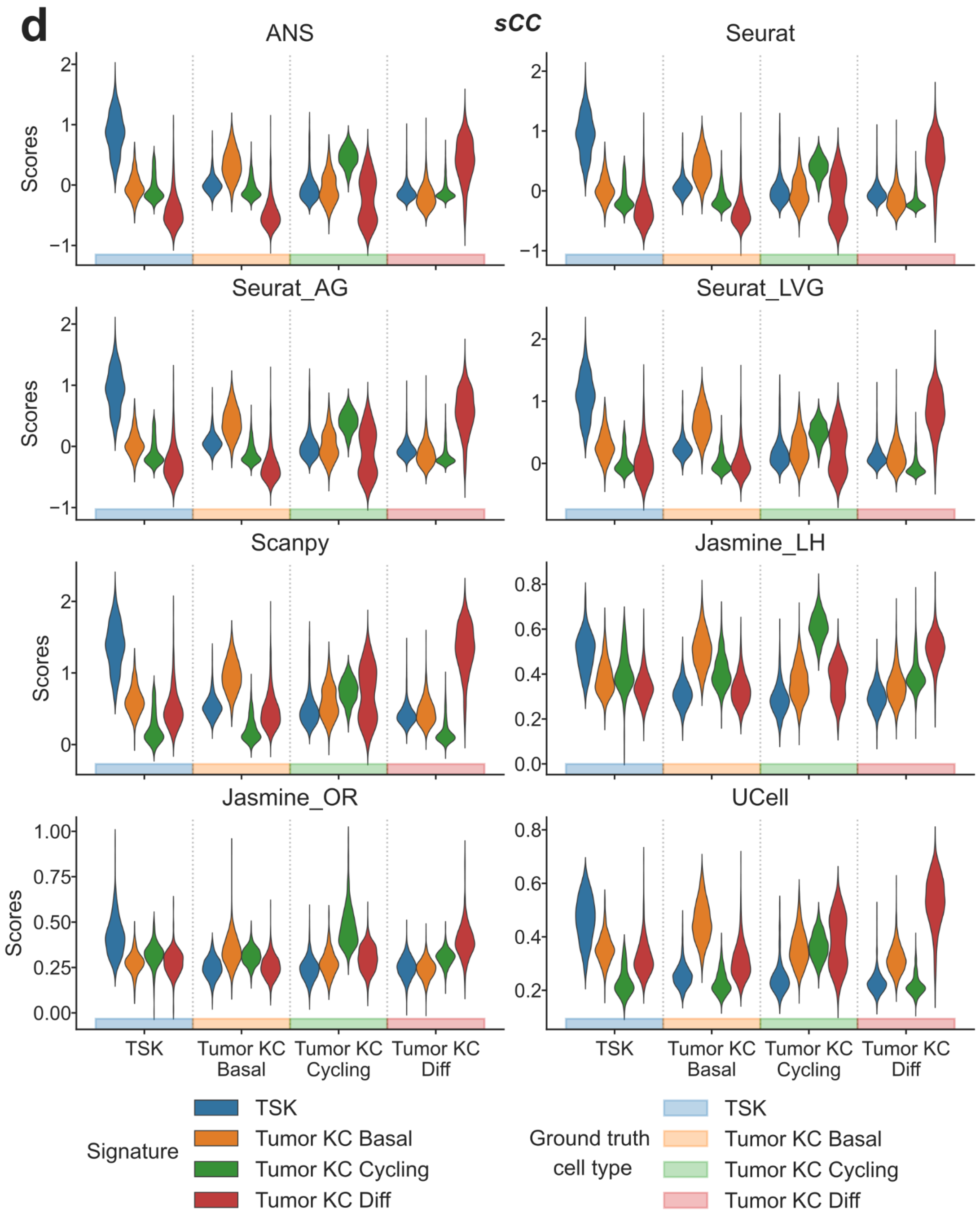

d
sCC
ANS
Seurat
Seurat_AG
Seurat_LVG
Scanpy
Jasmine_LH
Jasmine_OR
UCell
Scores
TSK
Tumor KC Basal
Tumor KC Cycling
Tumor KC Diff
Signature
Ground truth cell type

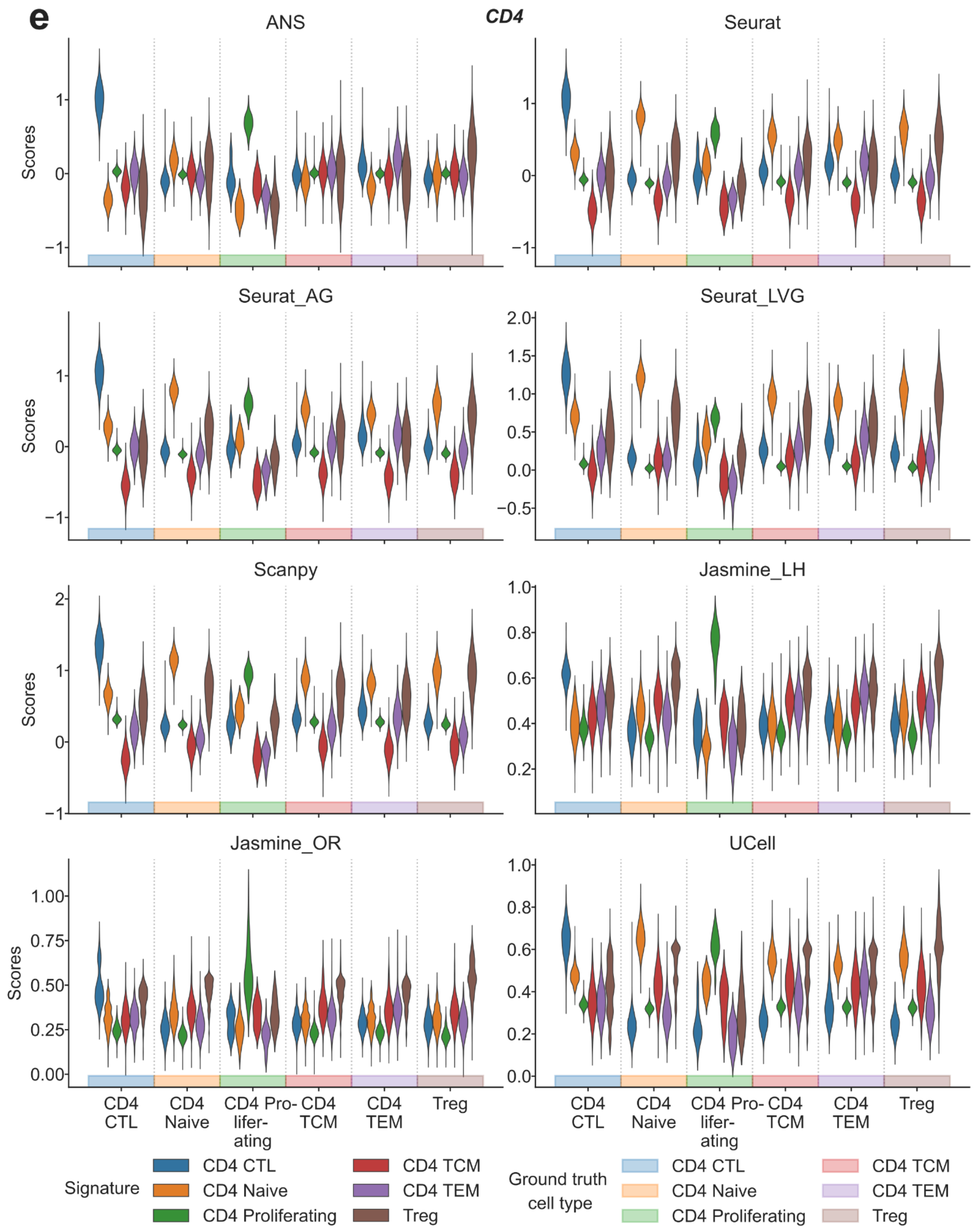

e
CD4
ANS
Seurat
Seurat_AG
Seurat_LVG
Scanpy
Jasmine_LH
Jasmine_OR
UCell
Scores
CD4 CTL
CD4 Naive
CD4 Pro-lifer-ating
CD4 TCM
CD4 TEM
Treg
Signature
CD4 CTL
CD4 Naive
CD4 Proliferating
CD4 TCM
CD4 TEM
Treg
Ground truth cell type
CD4 CTL
CD4 Naive
CD4 Proliferating
CD4 TCM
CD4 TEM
Treg

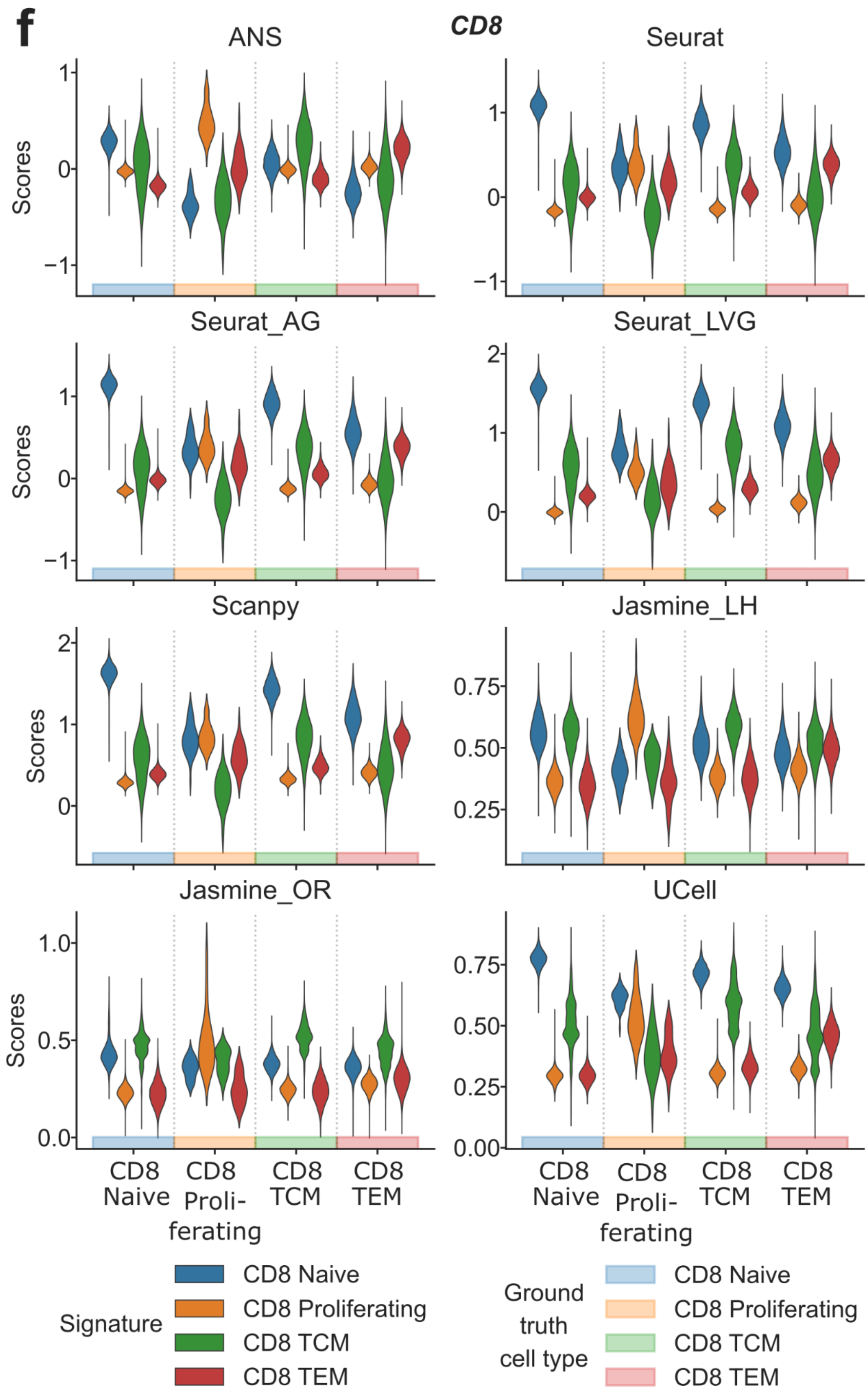

f
CD8
ANS
Seurat
Seurat_AG
Seurat_LVG
Scanpy
Jasmine_LH
Jasmine_OR
UCell
Scores
CD8 Naive
CD8 Proli-ferating
CD8 TCM
CD8 TEM
Signature
CD8 Naive
CD8 Proliferating
CD8 TCM
CD8 TEM
Ground truth cell type
CD8 Naive
CD8 Proliferating
CD8 TCM
CD8 TEM

**Figure S9:** Comparative analysis of scoring methods for cell type and state annotation. Original cell type and state signatures were used potentially containing overlapping gene sets. **a,** Score distributions for cell-type-specific signatures (B cells, monocytes, and NK cells) separated by true cell type annotations, calculated for each scoring method. **b**, Score distributions for B-cell subtype signatures separated by true cell subtypes, calculated by each scoring method. **c**, Row-normalized confusion matrix of B-cell subtype annotation based on the highest scores. **d**, Relationship between hard labelling performance and score information quantity in cancer vs. PBMC datasets. Scatterplots show balanced accuracy (x-axis) against score information quantity (y-axis) for various scoring method-dataset combinations. Balanced accuracy quantifies hard labelling performance, while score information quantity indicates the scores' effectiveness in subtype classification. The diagonal line indicates perfect metric alignment, with vertical distances from this line representing scale imbalance. **e**, Quantitative analysis of scale imbalance across scoring methods and tissue types (cancer vs. PBMC). The mean and standard deviation of scale imbalance for each method are shown. Scale imbalance is the absolute difference between score information quantity and balanced accuracy in direct label assignment. The method with the lowest mean scale imbalance, indicating optimal consistency between information content and labelling accuracy, is highlighted in bold. **f**, Cell-state and -type annotation performance overview for all eight datasets and scoring methods.

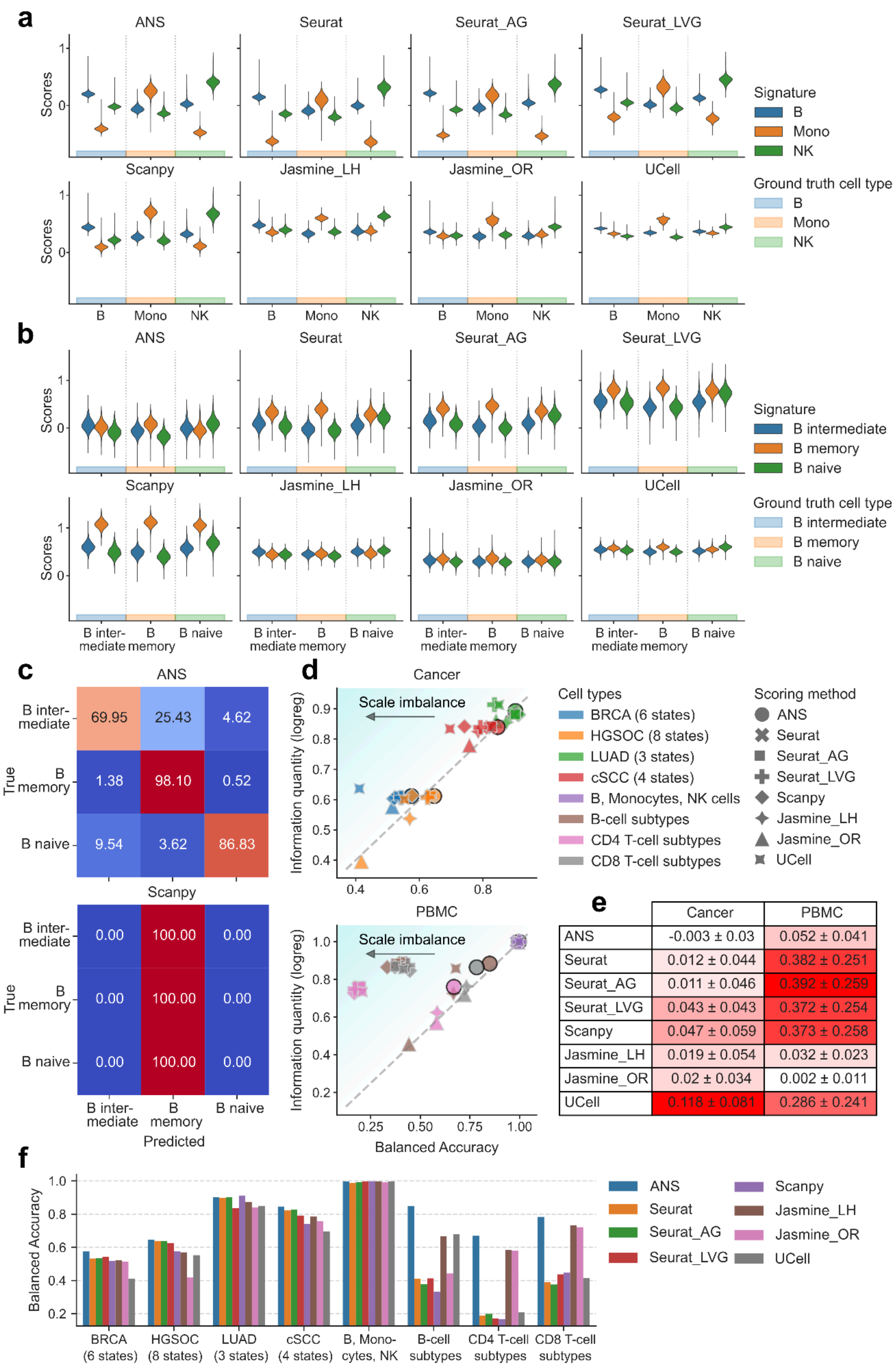


| | Cancer | PBMC |
|---|---|---|
| ANS | -0.003 ± 0.03 | 0.052 ± 0.041 |
| Seurat | 0.012 ± 0.044 | 0.382 ± 0.251 |
| Seurat_AG | 0.011 ± 0.046 | 0.392 ± 0.259 |
| Seurat_LVG | 0.043 ± 0.043 | 0.372 ± 0.254 |
| Scanpy | 0.047 ± 0.059 | 0.373 ± 0.258 |
| Jasmine_LH | 0.019 ± 0.054 | 0.032 ± 0.023 |
| Jasmine_OR | 0.02 ± 0.034 | 0.002 ± 0.011 |
| UCell | 0.118 ± 0.081 | 0.286 ± 0.241 |

**Figure S10:** Confusion matrices for cell type/state prediction across multiple methods and datasets. **a**, Comparison of prediction performance between ANS, Jasmine_LH, Jasmine_OR, and Scanpy for cancer datasets (BRCA, LUAD, HGSOC, cSCC). **b**, Results for Seurat, Seurat_AG, Seurat_LVG, and UCell on the same cancer datasets. **c-d**, Performance evaluation across immune cell subsets using the same methods, showing B/Monocytes/NK cells, B-cell subtypes, CD4 T-cell subtypes, and CD8 T-cell subtypes. Each matrix shows the relationship between true (y-axis) and predicted (x-axis) cell states, with values summing to one per row. The color scale ranges from blue (0) to red (1), with balanced accuracy (bal. acc.) shown for each matrix. Overlapping/ Original gene signatures were used to score cell states/types in each dataset.

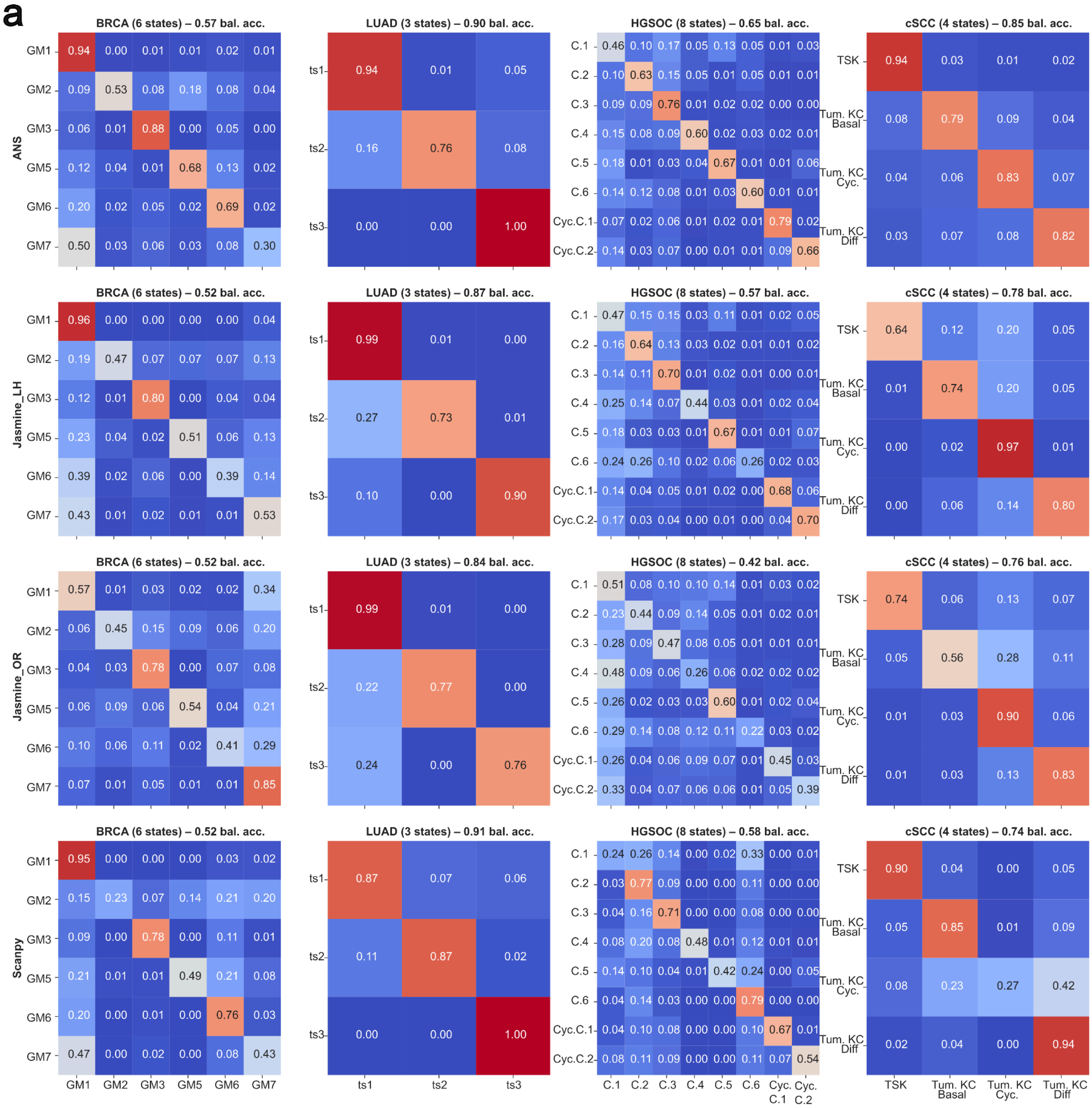

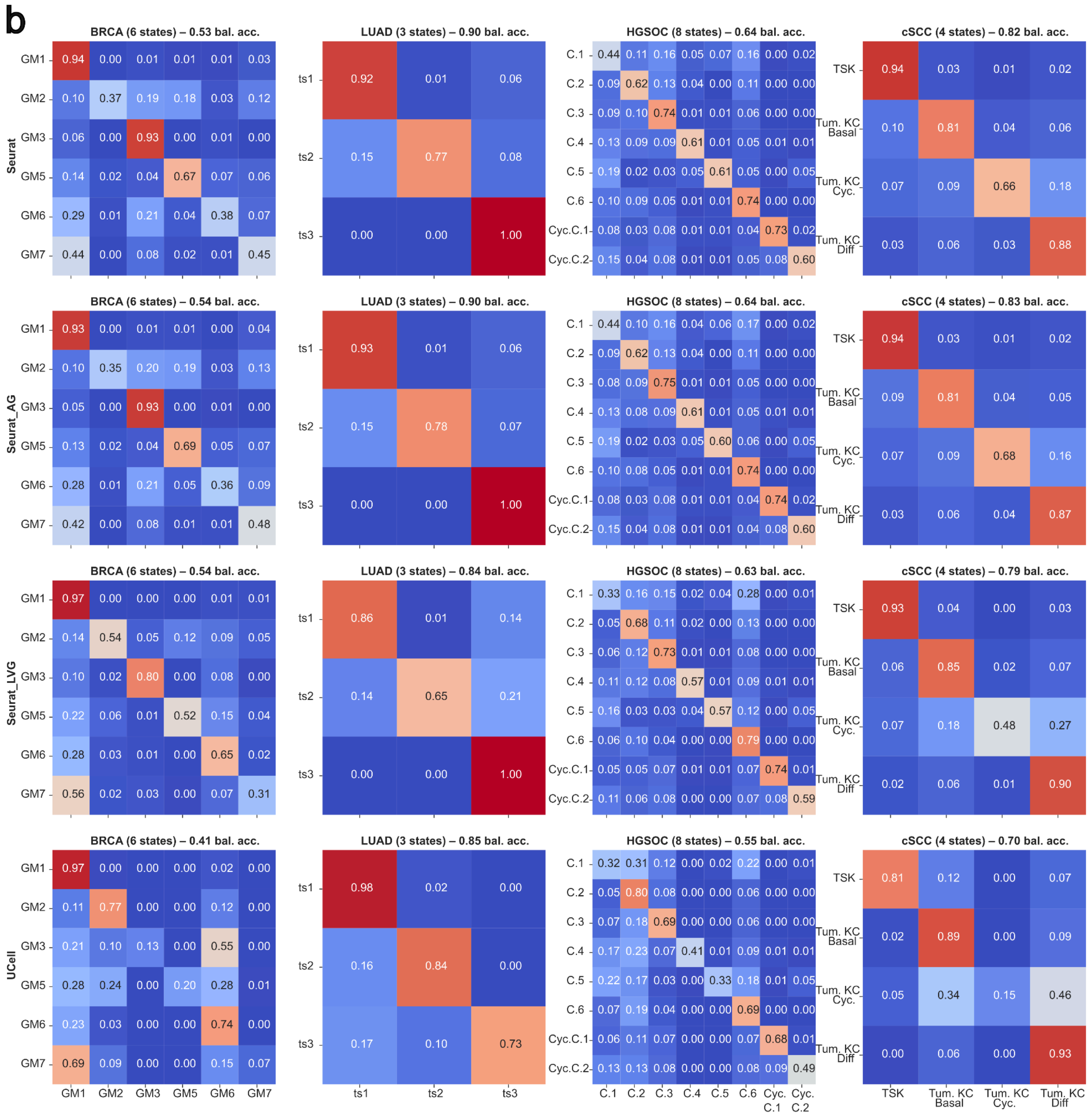
b
BRCA (6 states) – 0.53 bal. acc.
LUAD (3 states) – 0.90 bal. acc.
HGSOC (8 states) – 0.64 bal. acc.
cSCC (4 states) – 0.82 bal. acc.
BRCA (6 states) – 0.54 bal. acc.
LUAD (3 states) – 0.90 bal. acc.
HGSOC (8 states) – 0.64 bal. acc.
cSCC (4 states) – 0.83 bal. acc.
BRCA (6 states) – 0.54 bal. acc.
LUAD (3 states) – 0.84 bal. acc.
HGSOC (8 states) – 0.63 bal. acc.
cSCC (4 states) – 0.79 bal. acc.
BRCA (6 states) – 0.41 bal. acc.
LUAD (3 states) – 0.85 bal. acc.
HGSOC (8 states) – 0.55 bal. acc.
cSCC (4 states) – 0.70 bal. acc.
Seurat
Seurat_AG
Seurat_LVG
UCell
GM1 GM2 GM3 GM5 GM6 GM7
ts1 ts2 ts3
C.1 C.2 C.3 C.4 C.5 C.6 Cyc.C.1 Cyc.C.2
TSK
Tum. KC Basal
Tum. KC Cyc.
Tum. KC Diff

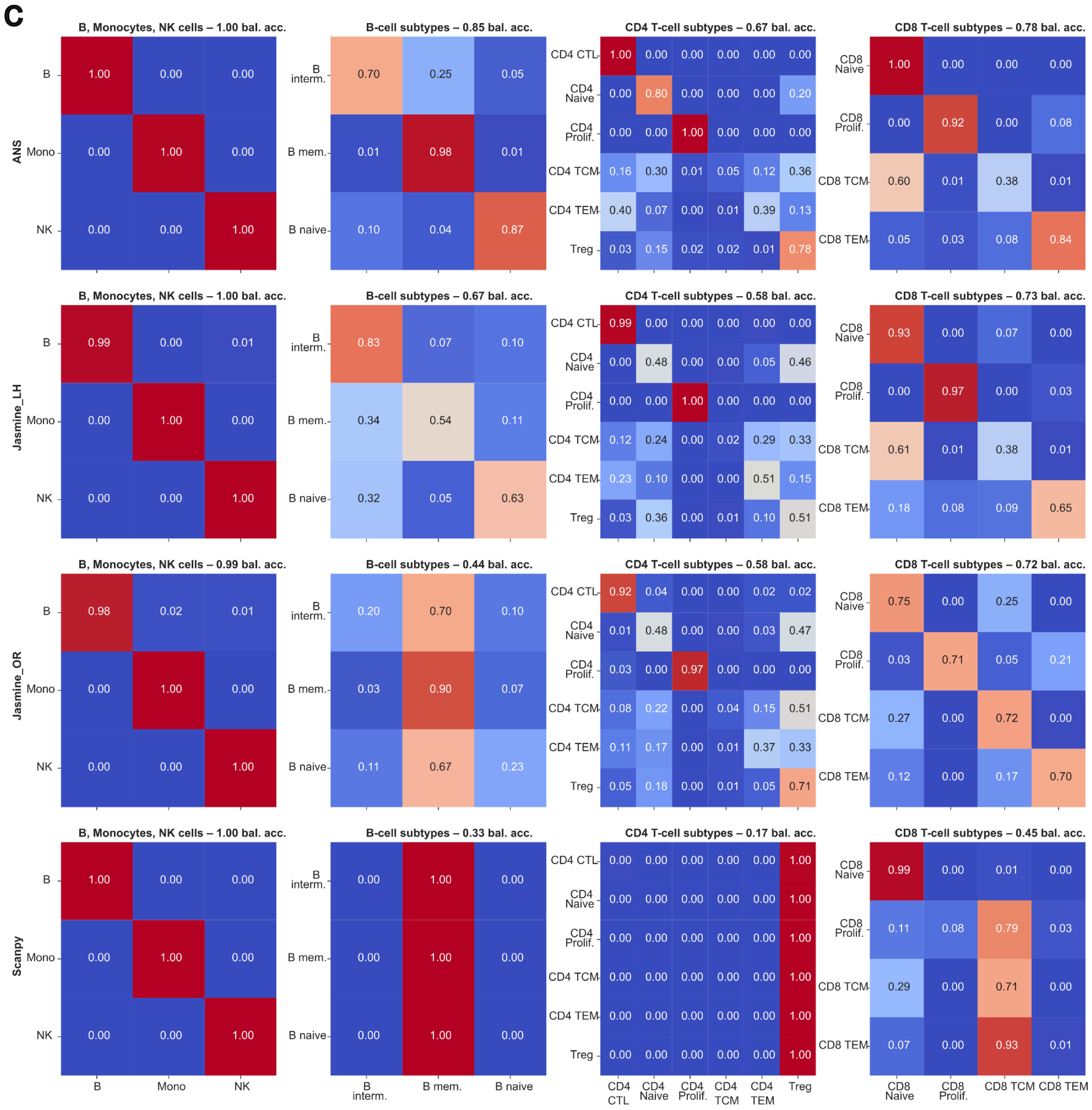
C
B, Monocytes, NK cells – 1.00 bal. acc.
B-cell subtypes – 0.85 bal. acc.
CD4 T-cell subtypes – 0.67 bal. acc.
CD8 T-cell subtypes – 0.78 bal. acc.
B, Monocytes, NK cells – 1.00 bal. acc.
B-cell subtypes – 0.67 bal. acc.
CD4 T-cell subtypes – 0.58 bal. acc.
CD8 T-cell subtypes – 0.73 bal. acc.
B, Monocytes, NK cells – 0.99 bal. acc.
B-cell subtypes – 0.44 bal. acc.
CD4 T-cell subtypes – 0.58 bal. acc.
CD8 T-cell subtypes – 0.72 bal. acc.
B, Monocytes, NK cells – 1.00 bal. acc.
B-cell subtypes – 0.33 bal. acc.
CD4 T-cell subtypes – 0.17 bal. acc.
CD8 T-cell subtypes – 0.45 bal. acc.
ANS
Jasmine_LH
Jasmine_OR
Scanpy
B
Mono
NK
B interm.
B mem.
B naive
CD4 CTL
CD4 Naive
CD4 Prolif.
CD4 TCM
CD4 TEM
Treg
CD8 Naive
CD8 Prolif.
CD8 TCM
CD8 TEM

d

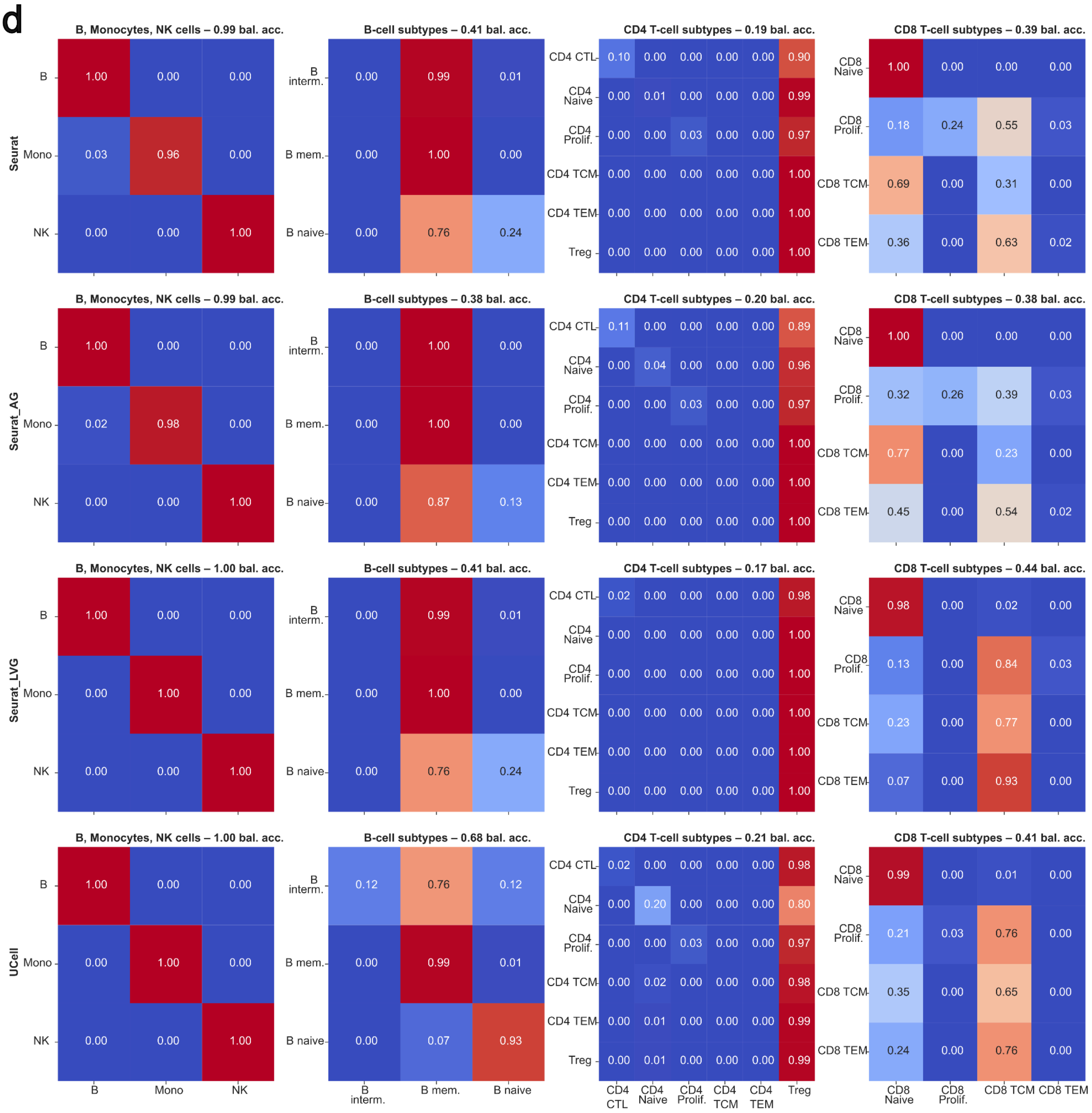

B, Monocytes, NK cells – 0.99 bal. acc.
B-cell subtypes – 0.41 bal. acc.
CD4 T-cell subtypes – 0.19 bal. acc.
CD8 T-cell subtypes – 0.39 bal. acc.
B, Monocytes, NK cells – 0.99 bal. acc.
B-cell subtypes – 0.38 bal. acc.
CD4 T-cell subtypes – 0.20 bal. acc.
CD8 T-cell subtypes – 0.38 bal. acc.
B, Monocytes, NK cells – 1.00 bal. acc.
B-cell subtypes – 0.41 bal. acc.
CD4 T-cell subtypes – 0.17 bal. acc.
CD8 T-cell subtypes – 0.44 bal. acc.
B, Monocytes, NK cells – 1.00 bal. acc.
B-cell subtypes – 0.68 bal. acc.
CD4 T-cell subtypes – 0.21 bal. acc.
CD8 T-cell subtypes – 0.41 bal. acc.
Seurat
Seurat_AG
Seurat_LVG
UCell
B
Mono
NK
B interm.
B mem.
B naive
CD4 CTL
CD4 Naive
CD4 Prolif.
CD4 TCM
CD4 TEM
Treg
CD8 Naive
CD8 Prolif.
CD8 TCM
CD8 TEM

**Figure S11:** Score distributions by cell types for the remaining five pan-cancer EMT signatures on ESCC, LUAD_xing, CRC, and BRCA.

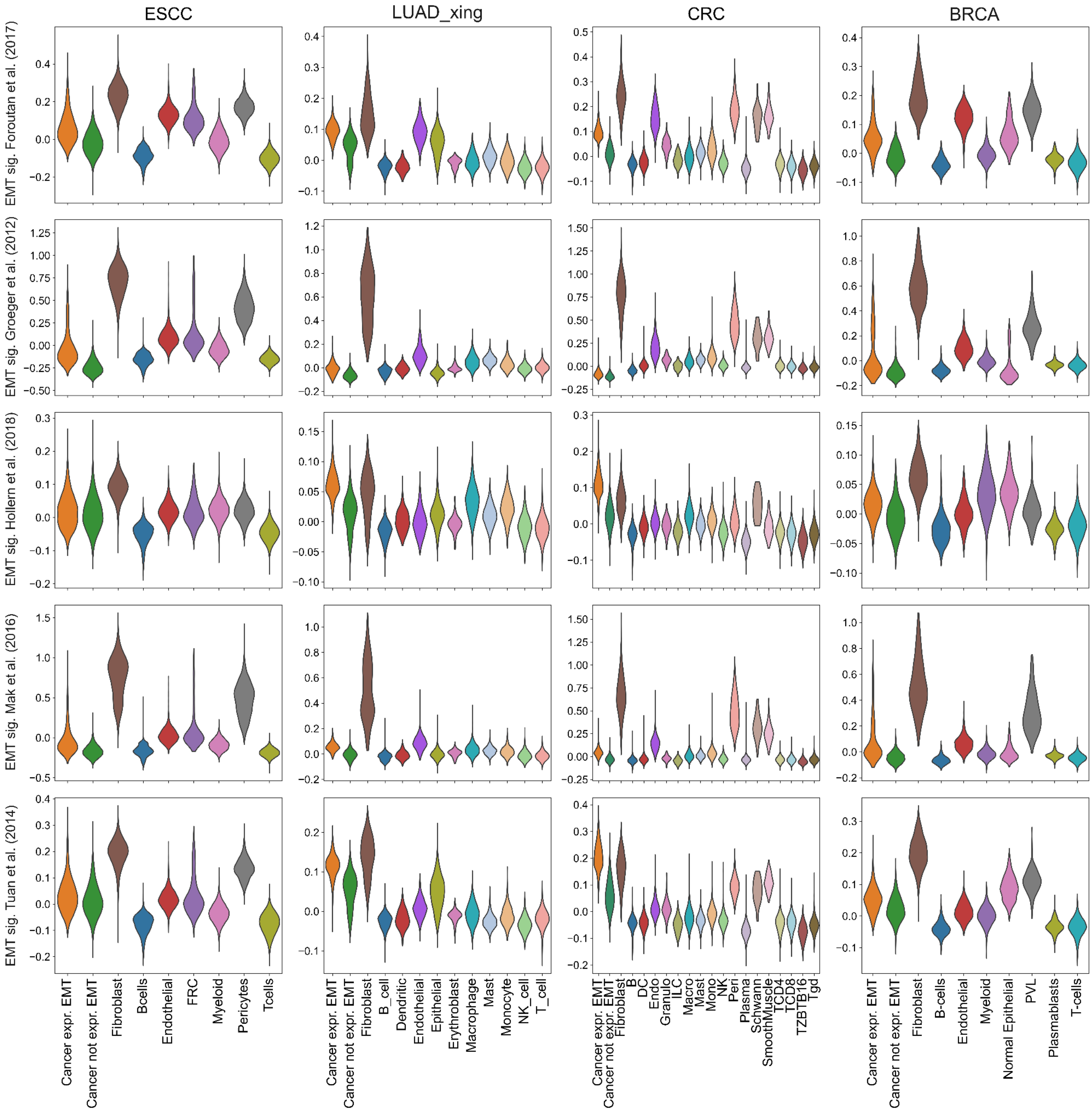

**Figure S12:** Dataset compositions for ESCC, LUAD, CRC, and breast carcinoma (BREAST), including cancer EMT cell annotations.

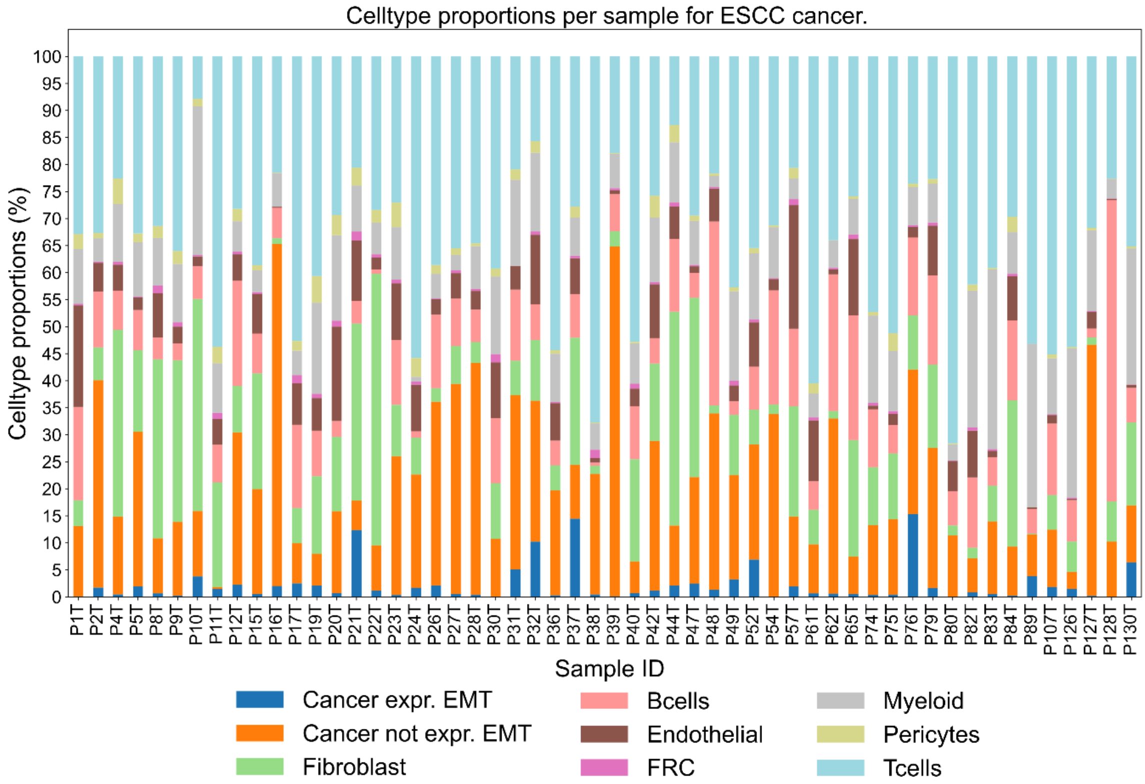


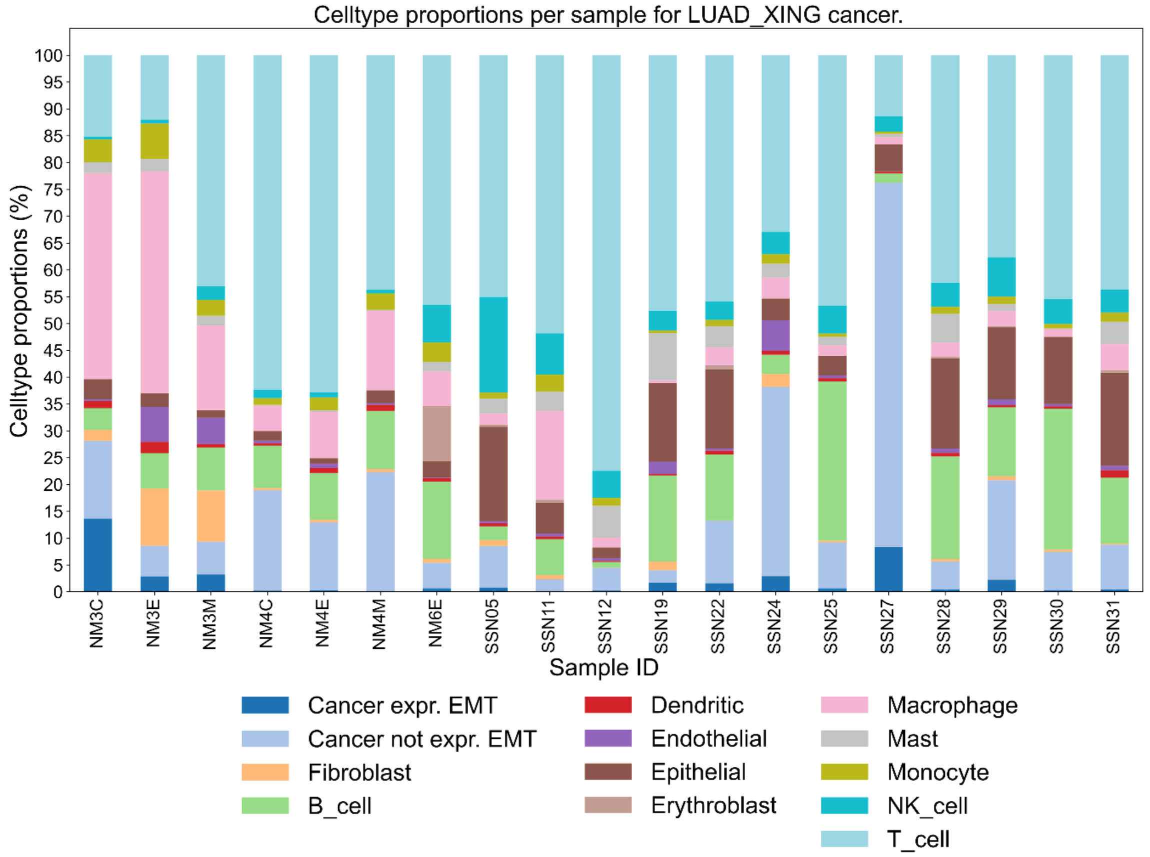

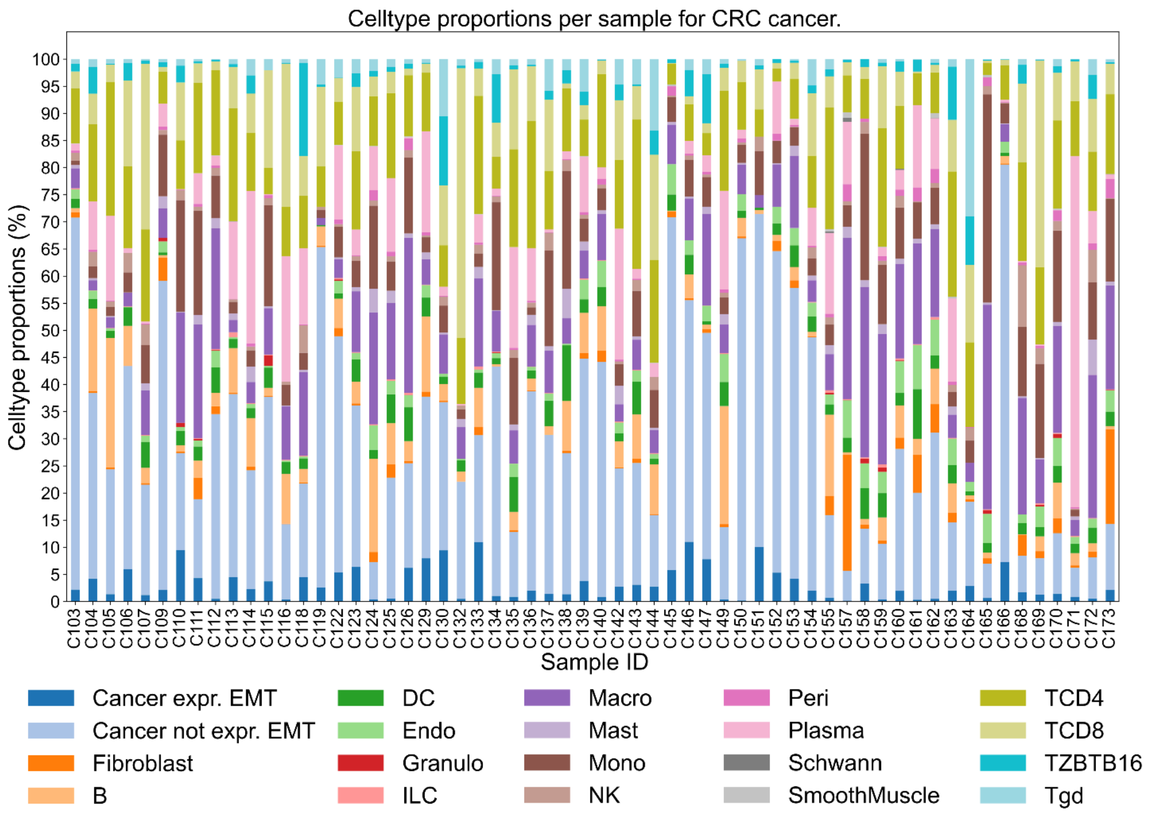
Celltype proportions per sample for CRC cancer.
Celltype proportions (%)
0 5 10 15 20 25 30 35 40 45 50 55 60 65 70 75 80 85 90 95 100
C103 C104 C105 C106 C107 C109 C110 C111 C112 C113 C114 C115 C116 C118 C119 C122 C123 C124 C125 C126 C129 C130 C132 C133 C134 C135 C136 C137 C138 C139 C140 C142 C143 C144 C145 C146 C147 C149 C150 C151 C152 C153 C154 C155 C157 C158 C159 C160 C161 C162 C163 C164 C165 C166 C168 C169 C170 C171 C172 C173
Sample ID
Cancer expr. EMT
Cancer not expr. EMT
Fibroblast
B
DC
Endo
Granulo
ILC
Macro
Mast
Mono
NK
Peri
Plasma
Schwann
SmoothMuscle
TCD4
TCD8
TZBTB16
Tgd

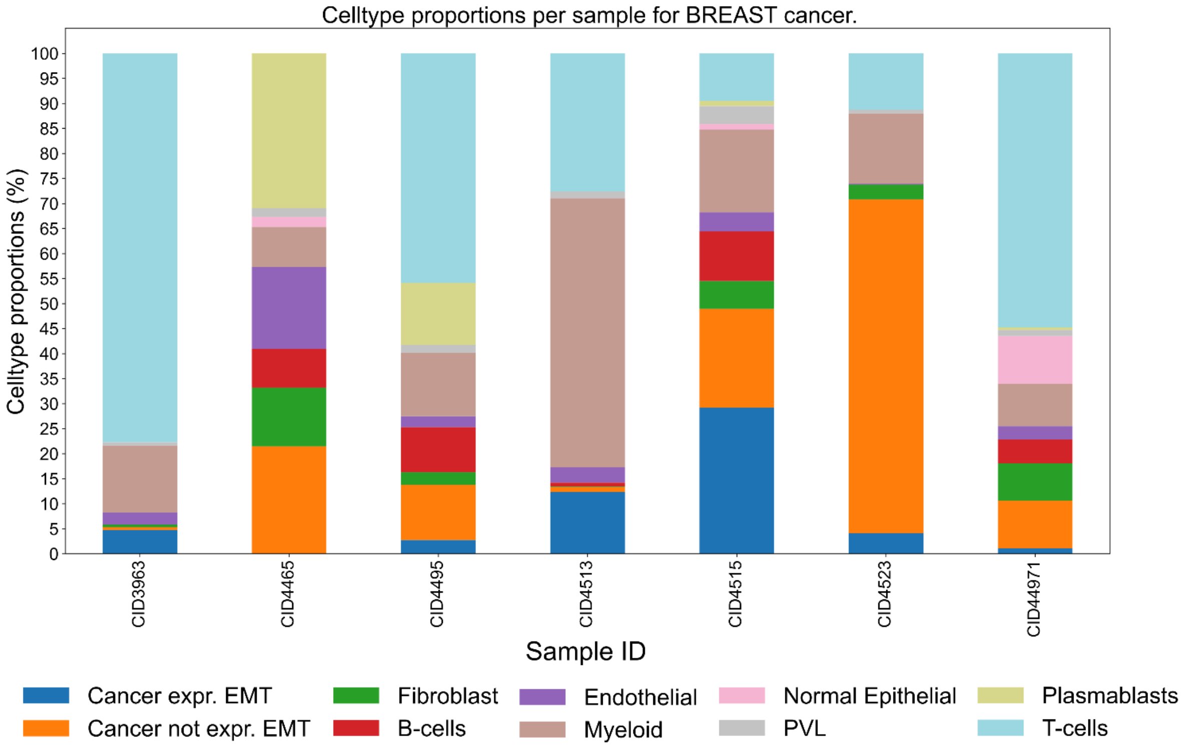
Celltype proportions per sample for BREAST cancer.
Celltype proportions (%)
0 5 10 15 20 25 30 35 40 45 50 55 60 65 70 75 80 85 90 95 100
CID3963 CID4465 CID4495 CID4513 CID4515 CID4523 CID44971
Sample ID
Cancer expr. EMT
Cancer not expr. EMT
Fibroblast
B-cells
Endothelial
Myeloid
Normal Epithelial
PVL
Plasmablasts
T-cells

**Figure S13**: Comparison of cell-state annotation performance in ovarian cancer using different data preprocessing approaches. The balanced accuracy of the considered scoring methods is shown for two data processing pipelines: (1) GEO data (GSE180661) with 1% gene filtering and mean-shift log-normalization on cancer cells (HGSOC 8 states), and (2) Raw data filtered using curated cells from cellxgene, followed by 1% gene filtering and mean-shift log-normalization (HGSOC 8 states, cellxgene)[1]. Results are presented for both original signatures and signatures without overlapping genes, with two different control gene selection strategies: excluding signature genes from the overall control gene pool (gene_pol=True) or allowing them to be included (gene_pol=False). The highly similar performance between both approaches suggests that the GEO dataset primarily contains the curated cells, and additional filtering steps do not significantly impact the annotation results. ANS consistently shows the highest balanced accuracy (~0.65) across all conditions, while Jasmine_OR performs the poorest (~0.40).

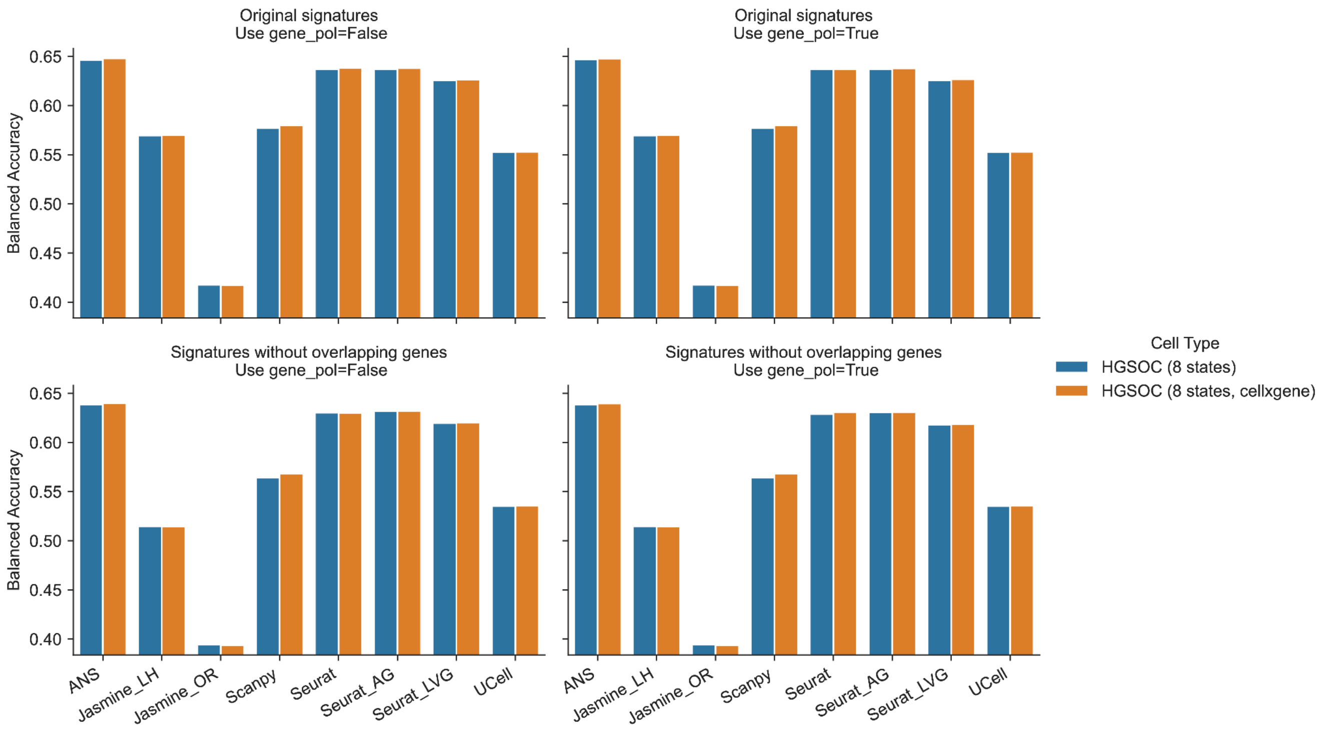


[1]CellxGene preprocessed dataset:
https://cellxgene.cziscience.com/collections/4796c91c-9d8f-4692-be43-347b1727f9d8

**Figure S14:** Comparison of cell-state annotation performance in cutaneous squamous cell carcinoma (cSCC) using different preprocessing approaches. The balanced accuracy of the considered scoring methods is shown for two data processing pipelines: (1) GEO data (GSE144240, GSE144236) with 1% gene filtering and mean-shift log-normalization on cancer cells (cSCC 4 statess, self-pp), and (2) Data processed through the CanSig pipeline followed by 1% gene filtering and mean-shift normalization (cSCC 4 states). Results are presented for both original signatures and signatures without overlapping genes, with two different control gene selection strategies: excluding signature genes from the overall control gene pool (gene_pol=True) or allowing them to be included (gene_pol=False). The similar performance patterns between both approaches indicate that the preprocessing steps in the CanSig pipeline do not significantly alter the annotation results. ANS demonstrates the highest balanced accuracy (~0.85) across all conditions, while UCell consistently shows the lowest performance (~0.70).

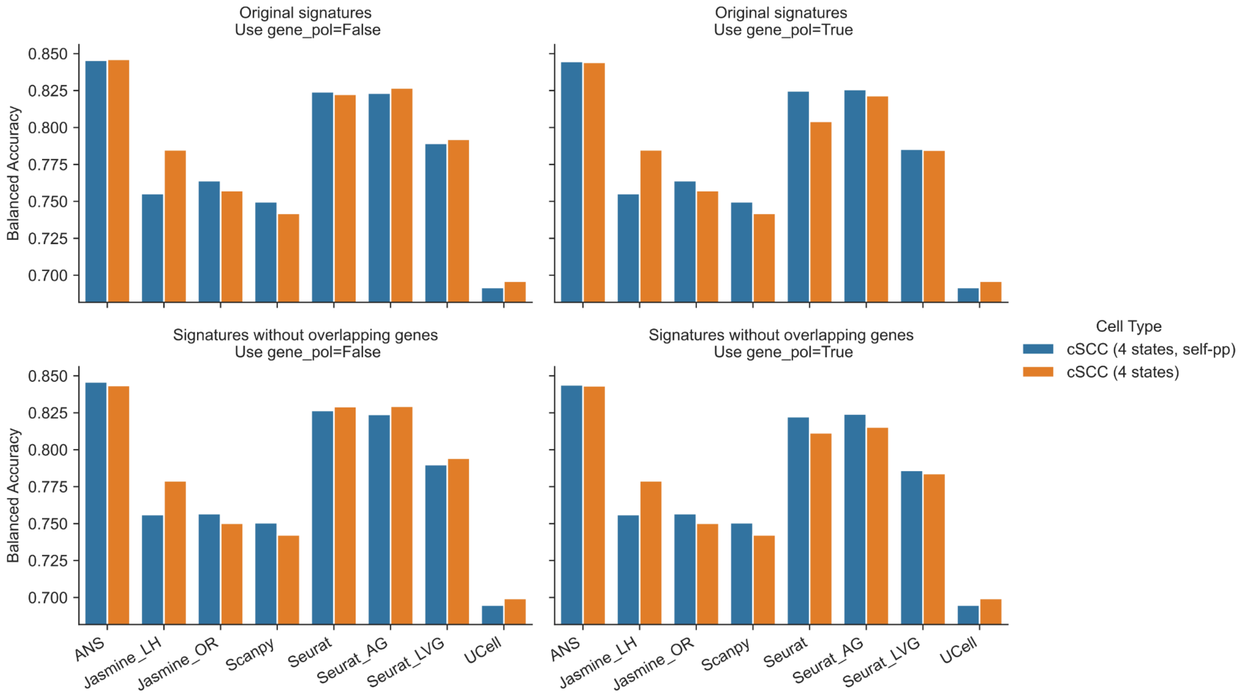

**Figure S15:** Impact of data preprocessing methods and gene selection for cell states annotation. We compared two preprocessing approaches: the CanSig pipeline and the 3CA pipeline[2], which implements different cell and gene filtering criteria. The ovarian cancer dataset was processed using 3CA only. For the lung adenocarcinoma (LUAD) dataset from Kim et al., we performed parallel analyses using both CanSig and 3CA preprocessing methods. In both cases, we applied additional filtering to remove genes expressed in less than 1% of cells and performed log-normalization before calculating signature scores. For methods requiring a gene pool for control gene selection, we evaluated two strategies: (1) excluding all signature genes from the pool and (2) placing no restrictions on pool composition. The figure displays a matrix where rows show the balanced accuracy for signatures with overlapping versus non-overlapping genes, and columns indicate whether a gene pool was utilized.

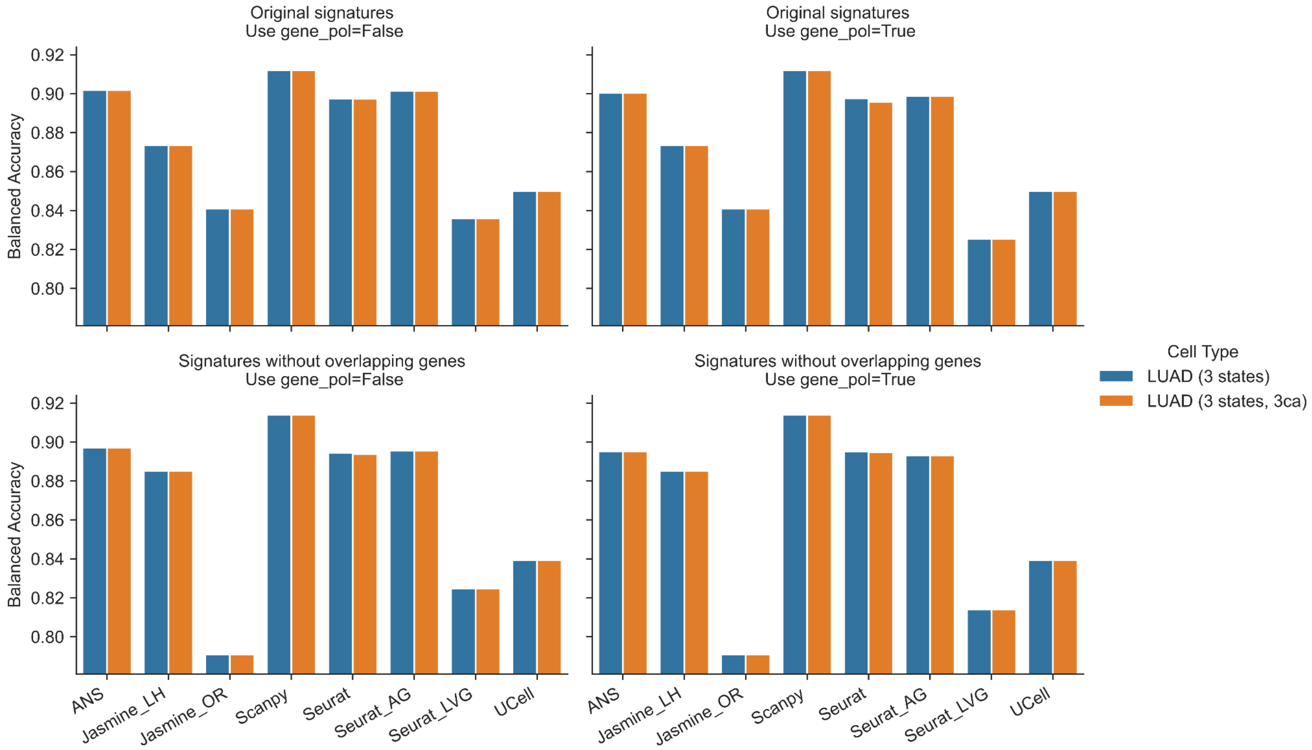


[2]3CA: Curated Cancer Cell Atlas (https://www.weizmann.ac.il/sites/3CA/) contains multiple curated cancer scRNA-seq datasets. 3CA uses its own pipeline to curate datasets.
